# The type I and III interferon responses restrict infection with tick-borne orthoflaviviruses through IFI6

**DOI:** 10.1101/2025.04.06.647422

**Authors:** Felix Streicher, Devin Kenney, Vincent Caval, Maxime Chazal, Sophie-Marie Aicher, Ségolène Gracias, Ferdinand Roesch, Florian Douam, Nolwenn Jouvenet

## Abstract

Tick-borne orthoflaviviruses (TBOVs) are a growing global health concern. Several representatives of this viral family cause fatal disease in humans with increasing case numbers throughout the last decades. The innate immune response, especially interferon (IFN)-dependent signaling, is an essential part of the human defense system that counteracts infection with TBOVs and other viruses. Even though they activate the same signaling cascade, IFNs belonging to the type I and III families trigger differing gene expression patterns. Which genes the two IFN families induce to restrict infection with TBOVs remains poorly characterized. Here we show that type I and III IFNs are both capable of restricting TBOV infection of human cell lines in a cell type-specific manner. Infection of C57BL/6J mice with knockouts for either IFN type I or III receptors further underscored the critical role of IFN signaling in controlling TBOV replication *in vivo*. To assess the contribution of single genes to controlling TBOV infection in human cells, we used a CRISPR/Cas9-KO-based screening approach. This strategy identified IFI6 as a central player for IFN type I- and III-driven responses against TBOVs. We further defined IFI6 as an ER-resident protein that restricts TBOV replication at a post-entry step. Our work thus opens new perspectives for targeting weak points in the life cycle of TBOVs and other orthoflaviviruses, potentially paving the way for the development of new antiviral therapeutics.

**One Sentence Summary:** Type I and III interferons are crucial for protection against tick-borne orthoflavivirus infection *in vitro* and *in vivo*, both relying on IFI6 as a main antiviral effector.

## INTRODUCTION

Tick-borne orthoflaviviruses (TBOVs) are enveloped viruses with a 10-11kb long, single-stranded positive-sense RNA genome encoding for three structural (Capsid (C), Membrane (M) and Envelope (E)) and 7 non-structural (NS1, NS2A, NS2B, NS3, NS4A, NS4B and NS5) proteins (1). Much like their more extensively studied mosquito-borne relatives, such as Zika virus (ZIKV), Dengue virus (DENV) or yellow fever virus (YFV), TBOVs pose a major threat to global health (1–4). Several TBOVs cause severe diseases in humans and can be divided into viruses that primarily induce hemorrhagic or neuronal pathology. Members of the latter, such as tick-borne encephalitis virus (TBEV), Powassan virus (POWV), or Louping-ill virus (LIV), infect the human central nervous system (CNS) and cause long-term neurological damage or death (3,5–7). They are transmitted by a variety of mostly hard-bodied ticks of predominantly *Ixodes* and *Dermacentor spp*. that are distributed worldwide with a geographical preference for the temperate zone of the northern hemisphere (2,3). Although most human infections occur through tick bites, numerous cases of TBOV infection after ingestion of unpasteurized milk stemming from infected livestock such as sheep and goats have been reported in humans (8,9). As of 2025, TBEV, the prototypical neuropathogenic TBOV that is responsible for the most severe infections in humans, is the only TBOV against which a vaccine was approved for use in humans. Despite this, the incidence of TBEV-induced cases of severe disease steeply increased throughout the last decades (10–12). This is linked to a constant expansion of the habitats of the vector tick species and low vaccination rates, but also results from the lack of antiviral treatment options against TBOVs (7,13,14). To counteract the continuous rise in the number of patients suffering from severe disease caused by these emerging pathogens, alternative strategies against TBOVs need to be explored.

The first line of defense against pathogenic intruders in eukaryotes is the innate immune system (15). The mammalian innate immune system has evolved to rapidly control virus replication by inducing the expression of antiviral cytokines like type I and III interferons (IFNs). Upon secretion by infected cells, IFN type I (IFNα, IFNβ) and type III (IFNλ, also known as IL29, IL28A and IL28B) bind to their heterodimeric receptors, IFNAR1/IFNAR2 and IFNλR1/IL-10R2, respectively, and activate the canonical JAK/STAT pathway in infected and surrounding cells (16). Activation of this pathway results in the expression of a plethora of IFN-stimulated genes (ISGs) that concertedly establish an antiviral state in the cells. ISGs comprise a core set of genes that, upon stimulation, are induced at high levels in all cell types across mammalian species (17). Certain ISGs directly block the viral life cycle by targeting specific stages of virus replication, including entry into host cells, protein translation, replication, or assembly of new viral particles (18). While some ISGs display virus-(or viral family-) specific antiviral activity, others exhibit broad-spectrum antiviral functions. Furthermore, ISGs are also involved in the regulation of IFN signaling and thus are key for facilitating the return to cellular homeostasis. However, the contribution of most ISGs to the modulation of viral infection and replication remains poorly characterized (18).

Despite both activating the JAK/STAT pathway, IFN-I and IFN-III stimulate ISGs with different kinetics and potency, and in a cell type-dependent manner (19–22). These differences are likely connected to the abundance of type I or III receptor complexes expressed at the surface of different cell types (21). While type I IFNs and their receptors are ubiquitously expressed and robust inducers of the antiviral state in most cell types, type III IFN receptor complexes are expressed in a limited set of cell types, mainly in cells that compose mucosal surface tissues, like the lung or the gut (23–26). The importance of IFN-I signaling in controlling TBOV infections was previously established both in several distinct human cell types *in vitro* and in various murine models *in vivo* (6,27–36). In contrast, the antiviral potential of type III IFNs against TBOVs remains unknown. In addition, although the antiviral activities of IFN-I against TBOVs are documented, their effectors are poorly characterized.

Aiming to clarify this, we investigated the impact of IFN-I and -III on TBOV replication *in vitro* and *in vivo* and performed the first CRISPR/Cas9 knock-out genetic screen for this viral family to identify which IFN-induced effector(s) contribute(s) to the anti-TBOV state in human cells.

## RESULTS

### Human cell lines derived from tissues with physiological relevance to TBOV pathogenesis are susceptible to infection with two TBOVs

The transmission of TBOVs to humans can occur either through tick bite or ingestion of unpasteurized milk from infected animals (3,7,9). The latter marks cells of the intestinal epithelium as potential first targets of TBOV infections following oral transmission, which has been sparsely investigated to date (37). Independent of the transmission route, viral replication in neurons constitutes an essential part of the pathogenesis of neurotropic TBOVs (38). Thus, we selected human colorectal adenocarcinoma (Caco2) and medulloblastoma (DAOY) cells as human cell lines to study TBOV replication, since both were previously reported to be susceptible to TBEV (37,30). Caco2 and DAOY cells were infected with TBEV (strain Hypr) at a multiplicity of infection (MOI) of 0.1. Viral replication was monitored over time by analyzing the expression of the viral protein NS5 through immunofluorescence imaging at 24 and 48 h post-infection (hpi) (Fig.1A). Both cell lines were susceptible to TBEV, with an increase in NS5-positive cells over time (Fig.1A). In line with this, flow cytometric analysis using pan-orthoflavivirus anti-E antibodies revealed that while about 15% of Caco2 and 10% of DAOY cells were infected at 24 hpi, this increased to 55% and 45%, respectively, at 48 hpi (Fig.1B). Consistently, assessment of viral replication by RT-qPCR analysis showed significant increase of viral RNA over time (Fig.1C), further indicating robust TBEV replication in both cell lines. To assess whether these two cell lines were susceptible to other members of the TBOV group, they were infected with POWV (strain LB) at an MOI of 5. About 3 to 6 % of Caco2 and DAOY cells were positive for viral E proteins at 48 hpi (Fig.1D). Viral RNA yields increased over time (Fig.1E), further indicating viral replication. However, viral RNA abundances were about two logs lower in POWV-infected cells than in TBEV-infected cells (Fig.1C and E). In sum, despite disparities, Caco2 and DAOY cells were susceptible to TBEV and POWV. This validates these cell lines as useful models approximating tissues with physiological relevance to TBOV pathogenesis.

**Fig. 1.**
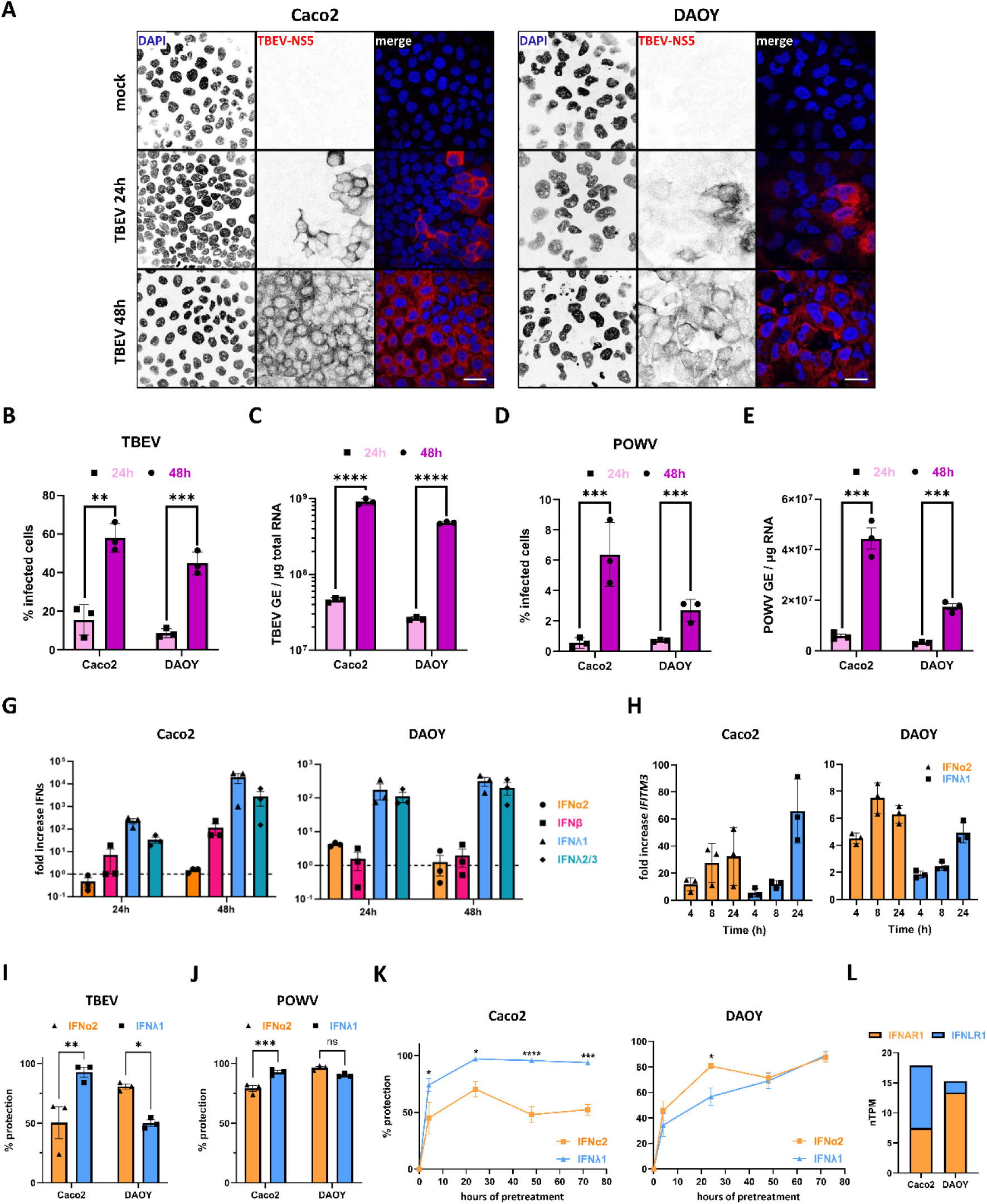
Type I and III IFNs protect human cell lines with physiological relevance against infection with TBOVs in a cell type-specific manner. **(A)** Confocal microscopy analysis of human colorectal adenocarcinoma (Caco2) or medulloblastoma cells (DAOY) infected with TBEV-Hypr at an MOI of 0.1. Twenty-four and forty-eight hours post-infection, cells were fixed, stained for TBEV-NS5 protein (red), and their nuclei stained with DAPI (blue). Scale bar 30 µm. Images are representative of two independent experiments. **(B-E)** Caco2 or DAOY cells were infected with either TBEV-Hypr (MOI 0.1) or POWV-LB (MOI 5) and monitored for percentage of infected cells *via* cytometric analysis after staining for orthoflaviviral E protein **(B, D)** or for relative amounts of viral RNA, expressed as genome equivalents (GE), *via* RT-qPCR analysis **(C, E)** at 24 and 48 hpi. **(F)** RT-qPCR analysis of mRNA levels of different interferons in Caco2 or DAOY cells infected with TBEV-Hypr at an MOI 1 for 24 or 48 h. **(G)** Caco2 or DAOY cells were treated with either recombinant human IFNα2 (50 U/mL, orange) or IFNλ1 (20 ng/mL, blue) for 16 h and assessed for *IFITM3* expression *via* RT-qPCR. **(H, I)** Cytometry-based analysis of infection levels in Caco2 and DAOY cells treated with either IFNα2 (50 U/mL, orange) or IFNλ1 (20 ng/mL, blue) for 16 h and infected with either TBEV-Hypr (MOI 1, **I**) or POWV-LB (MOI 5, J) for 24h. Cells were stained for orthoflaviviral envelope protein. **(K)** Caco2 or DAOY cells were treated with IFNα2 (50 u/mL, orange) or IFNλ1 (20 ng/mL, blue) for 4, 24, 48, or 72 h and infected with TBEV-Hypr (MOI 1) for 24 h before being stained for orthoflaviviral envelope protein and subjected to cytometry-based analysis for infection levels. **(L)** The abundance of IFNAR1 and IFNLR1 in cells of the Caco2 and DAOY cell lines as archived in the human protein atlas database (https://www.proteinatlas.org/ENSG00000142166-IFNAR1/cell+line; https://www.proteinatlas.org/ENSG00000185436-IFNLR1/cell+line) displayed as nTPM (transcripts per million). Data **(C-K)** from three independent experiments ± SEM. ns = non-significant, *P < 0.05, **P < 0.01, ***P < 0.001, ****P < 0.0001 by unpaired, two-tailed t-test **(C-F)** or two-way ANOVA with Sidak post-test **(I-K)**.

### Type I and III IFNs protect human cell lines against infection with TBOVs in a cell type-specific manner

Type I IFNs were previously shown to have a protective effect against several TBOVs *in vitro* (6,27,30,34). In contrast, the antiviral potential of the type III IFN family against TBOVs remains poorly characterized. We examined mRNA abundance of four IFN genes (*IFNA2*, *IFNB*, *IFNL1,* and *IFNL2/3)* in TBEV-infected cells at 24 and 48 hpi. Type III IFN upregulation surpassed type I IFN upregulation in TBEV-infected Caco2 and DAOY cells at both time points (Fig. 1G), potentially indicating an important role for type III IFNs in the antiviral response against TBOVs. To investigate whether Caco2 and DAOY cells respond differently to the two families of IFNs, the cells were treated with recombinant human IFNα2 or IFNλ1. Examination of the mRNA levels of *IFITM3*, a conserved ISG known to block orthoflaviviral replication (17,39,40), revealed that the highest induction was achieved with IFNλ1 treatment in Caco2 cells, whereas expression was more rapid but of lower magnitude after IFNα2 treatment in DAOY cells (Fig. 1H). To assess whether these differences translate into different levels of protection in Caco2 and DAOY cells, both cell lines were infected with TBEV after 16 hours of IFNα2 or IFNλ1treatment and the percentage of cells positive for viral E protein was assessed by flow cytometric analysis (Fig.1I). In line with stronger *IFITM3*-induction by IFN-III than IFN-I (Fig. 1H), Caco2 cells were protected against infection with TBEV following IFNλ1 treatment to around 90% compared to IFNα2 treatment, which protected only 50% of the cells from infection (Fig.1I). By contrast, DAOY cells mounted a stronger antiviral defense after exposure to IFNα2 compared to IFNλ1, correlating with the enhanced *IFITM3* induction by IFN-I previously observed in this cell line (Fig.1H). Similar trends of IFN protection were observed when cells were infected with POWV (Fig.1J). However, since POWV replicated at lower levels as compared to TBEV (Fig.1B-E), the differences in infection phenotype between IFN-treated cells and non-treated ones were more subtle. Due to the variations in the kinetics of the ISG response following stimulation by the distinct IFN families in the two cellular models (Fig.1K), we also examined the protective effect of IFNα2 and IFNλ1 against TBEV infection after various periods of pre-treatment. Caco2 cells were better protected by IFNλ1 than IFNα2 pre-treatment against infection with TBEV, independently of the timing (Fig.1K). At earlier times post-treatment, DAOY cells were better protected by IFNα2 than by IFNλ1 (Fig.1K). However, after extended incubation times, the two treatments triggered the same protective state (Fig.1K). To assess whether the varying degrees of protection were linked to the expression of the IFN receptor complexes in the two cell types, transcriptomic data from the human protein atlas were exploited (45). While Caco2 cells present a higher relative expression of *IFNLR1*, as compared to DAOY cells, DAOY cells express comparatively more *IFNAR1* than Caco2 cells (Fig.1L). This suggests that the abundance of receptors indeed influences the responsiveness to different IFN types and subsequently, the ability to control viral replication.

Altogether, this data shows that both type I and III IFNs mount an effective antiviral state against TBEV and POWV in Caco2 and DAOY cells with cell type-specific differences in potency of the antiviral response, which is likely related to the expression level of IFN receptors.

### IFN responses delay POWV-induced death in C57BL/6J mice

While few studies exploring the role of type I IFN signaling in TBOV infection in mice were conducted (29,37,41), it remains unknown to what extent type III IFNs are involved in the antiviral defense against TBOVs *in vivo*. We aimed to determine whether a lack of IFN-I signaling alone, or a lack of both IFN-I and IFN-III signaling, influences TBOV disease outcome and viral dissemination in mice. To do so, C57BL/6J mice genotyped as either WT, *Ifnar1^-/-^*, or *Ifnar1^-/-^ Ifnlr1^-/-^* were infected with POWV (10^3^ FFU) *via* subcutaneous injection into the footpad (Fig.2A, B). The survival rates of the infected mice were monitored for a 17-day period (Fig.2A), and viral titers in the sera were estimated through plaque assay at 2 dpi (Fig.2B). PBS-injected control mice survived the complete experiment without developing any clinical signs, independent of their genotype (Fig.2A). The majority of WT C57BL/6J exhibited an 80% mortality rate after 16 days of infection (Fig.2A). All *Ifnar1^-/-^* mice rapidly succumbed to infection, some as early as 3 dpi, and at the latest at 4 dpi (Fig.2A). Similarly, *Ifnar1^-/-^ Ifnlr1^-/-^* mice died four days after injection of POWV (Fig.2A). Titration of infectious POWV particles in the sera of *Ifnar1^-/-^* or *Ifnar1^-/-^ Ifnlr1^-/-^* mice two days after footpad injection yielded viral titers with no apparent difference between the two murine genotypes (Fig.2B). Therefore, following subcutaneous infection, *Ifnar1^-/-^* and *Ifnar1^-/-^ Ifnlr1^-/-^* mice were both hypersusceptible to POWV but exhibit no difference in terms of probability of survival or systemic viral titer. This suggests that IFN-I signaling supersedes IFN-III signaling in controlling POWV infection upon subcutaneous injection.

**Fig. 2.**
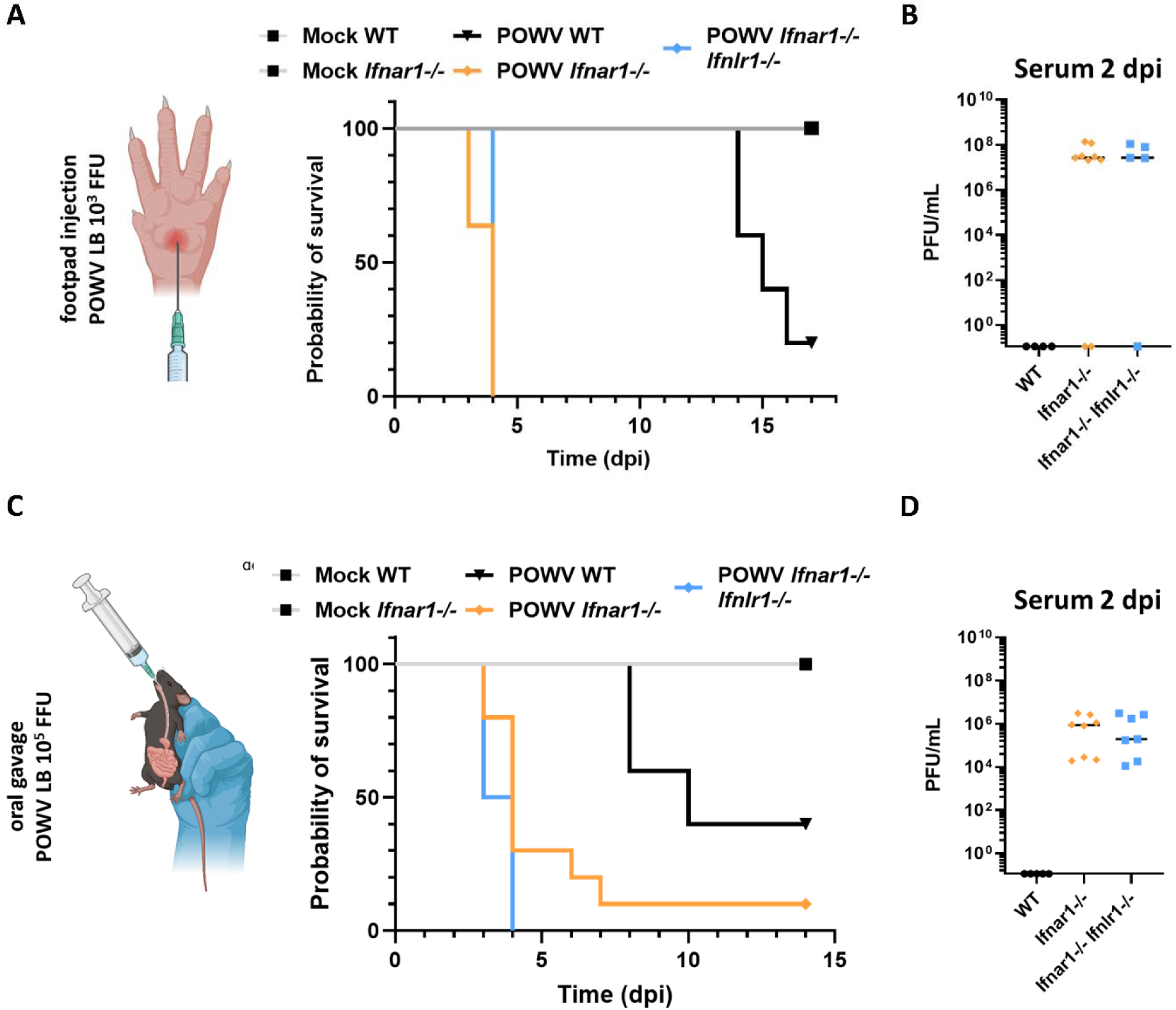
*Ifnar1^-/-^* and *Ifnar1^-/-^ Ifnlr1^-/-^* mice are hypersusceptible to POWV infection. **(A)** Survival of WT (black), *Ifnar1^-/-^* (orange), and *Ifnar1^-/-^ Ifnlr1^-/-^* (blue) C57BL/6J mice (5-10 per group) following subcutaneous POWV-LB infection with 10^3^ FFU or WT and *Ifnar1^-/-^* (grey, 2-3 per group) after administration of PBS as a control. **(B)** Titration of infectious viral particles in the sera of WT (black), *Ifnar1^-/-^* (orange), and *Ifnar1^-/-^ Ifnlr1^-/-^* (blue) C57BL/6J mice subcutaneously infected with 10^3^ FFU of POWV-LB (5-9 per group), as determined 2 dpi *via* plaque assay. **(C)** Survival of WT (black), *Ifnar1^-/-^* (orange), and *Ifnar1-/- Ifnlr1-/-* (blue) C57BL/6J mice (5-9 per group) following POWV-LB infection with 10^5^ FFU through oral gavage or WT and *Ifnar1^-/-^* (grey, 2-3 per group) after administration of PBS as a control. **(D)** Titration of infectious viral particles in the sera of WT (black), *Ifnar1^-/-^*(orange), and *Ifnar1^-/-^ Ifnlr1^-/-^* (blue) C57BL/6J mice orally infected with 10^5^ FFU of POWV-LB (4-8 per group), as determined 2 dpi *via* plaque assay. The figure was created using Biorender.com.

Our *in vitro* experiments using the Caco2 cell line suggest that cells of epithelial barrier tissues may respond more robustly to IFN-III treatment than IFN-I (Fig.1H). To test whether this translates to a crucial role of type III IFN-signaling in infection with TBOVs through ingestion, WT, *Ifnar1^-/-^*, or *Ifnar1^-/-^ Ifnlr1^-/-^* C57BL/6J mice were infected with 10^5^ infectious particles of POWV administered through oral gavage (Fig.2C). Mice were either monitored for survival for 14 days (Fig.2C) or sacrificed at 2 dpi and their sera examined for viral titers (Fig.2D). Mock-treatment did not affect any mice throughout the monitored period, whereas 60% of WT mice infected with POWV deceased within 10 days (Fig.2D). All *Ifnar1^-/-^ Ifnlr1^-/-^* mice succumbed to the virus in the first four days after infection, while 30% of *Ifnar1^-/-^* mice were still alive at this timepoint (Fig.2C). The *Ifnar1^-/-^* mice that were still alive at 4 dpi, except for one that survived the complete period of monitoring, died within the next three days. This delay in lethality might indicate a more prominent role of IFN-III, although still secondary to IFN-I signaling in the oral infection route of TBOVs. However, viremia in the sera was comparable between *Ifnar1^-/-^*and *Ifnar1^-/-^ Ifnlr1^-/-^* mice at 2 dpi, although the detected titers were 2-3 logs lower than in subcutaneously infected mice at the same timepoint (Fig.2B, D).

Together, these results indicate that type I IFN is the central player of the antiviral response against POWV *in vivo* and that while type III IFNs might support the inhibition of infection at mucosal surfaces like the intestine, this IFN signaling pathway does not appear critical to prevent rapid viremia and accelerated death in C57BL/6J mice.

### IFN responses prevent dissemination of POWV in a wide array of tissues in C57BL/6J mice

To characterize the effect of IFN-signaling on POWV dissemination and tropism in mice, WT, *Ifnar1^-/-^*, or *Ifnar1^-/-^ Ifnlr1^-/-^* C57BL/6J mice were infected either by subcutaneous injection into the footpad (10^3^ FFU, Fig.3A), or *via* oral gavage (10^5^ FFU, Fig.3B) and 2 dpi. Viral RNA abundance in individual tissues was quantified by RT-qPCR analysis. At 2 dpi, all the tested tissues of all mock-treated and WT mice subcutaneously infected with POWV were negative for viral RNA (Fig.3A), indicating that POWV replicated less efficiently in WT mice. By contrast, viral RNA was detected in all tissues of *Ifnar1^-/-^* and *Ifnar1^-/-^ Ifnlr1^-/-^* mice. Detection of viral RNA in the brain of *Ifnar1^-/-^*and *Ifnar1^-/-^ Ifnlr1^-/-^* mice, but not of WT mice, confirmed the importance of IFN-signaling for restriction of early dissemination to the brain (Fig. 3A). The highest levels of viral RNA were present in the spleen, suggesting that this tissue might be particularly dependent on IFN-signaling to limit viral dissemination. No significant difference in viral RNA load was observed between the two genotypes (Fig. 3A), indicating a widespread dissemination and a broad tissue-tropism of POWV in C57BL/6J mice lacking functional IFN signaling.

**Fig. 3.**
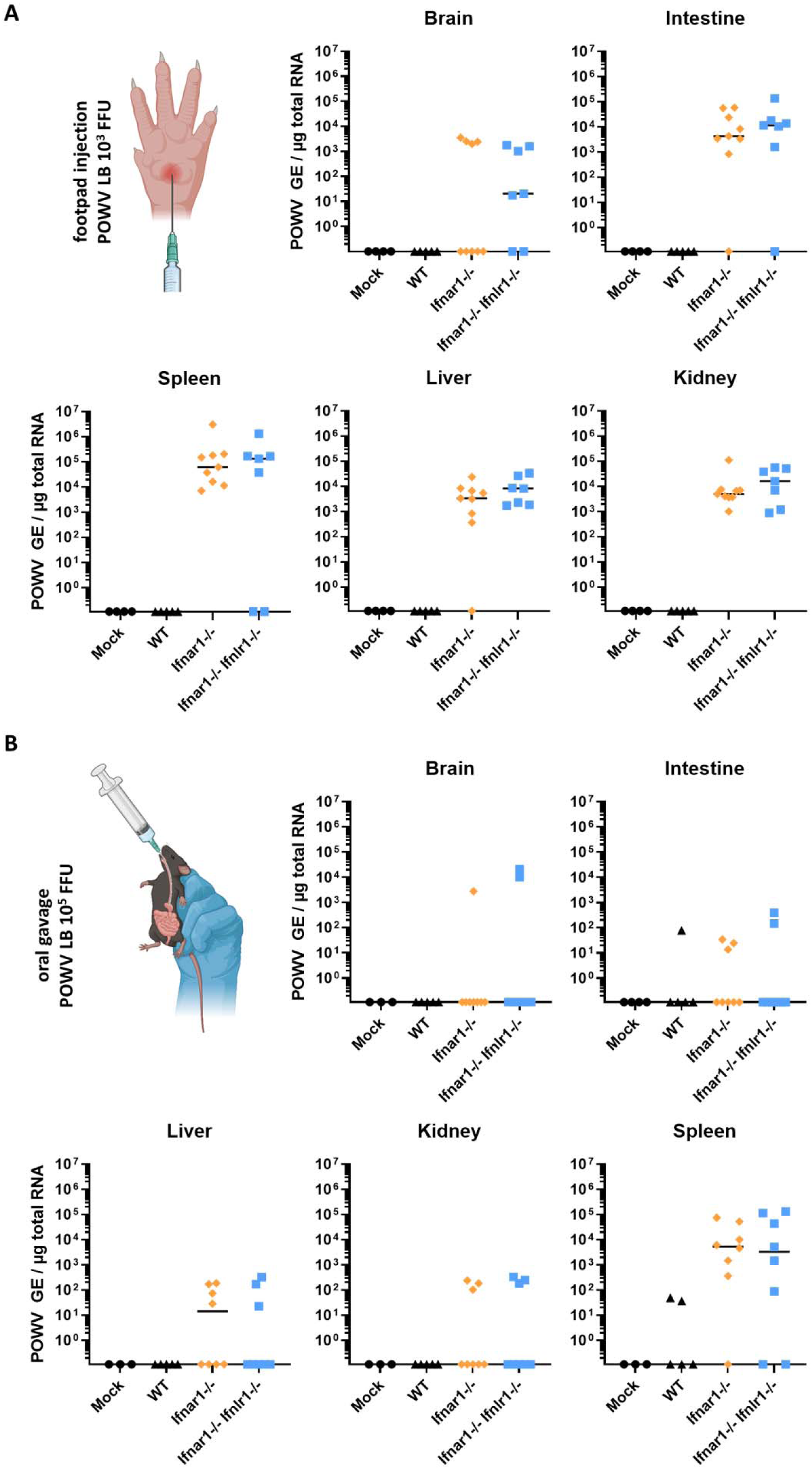
POWV shows widespread tropism in *Ifnar1^-/-^* and *Ifnar1^-/-^ Ifnlr1^-/-^* mice after injection in the footpad or administration of virus through oral gavage. **(A)** Viral loads within brain, small intestine, liver, kidney, and spleen tissue of WT (black), *Ifnar1^-/-^* (orange), and *Ifnar1^-/-^ Ifnlr^-/-^* (blue) C57BL/6J mice (4-9 per group) subcutaneously infected with 10^3^ FFU of POWV-LB, or **(B)** of WT (black), *Ifnar1^-/-^* (orange), and *Ifnar1^-/-^ Ifnlr1^-/-^* (blue) C57BL/6J mice (3-8 per group) infected with 10^5^ FFU of POWV-LB through oral gavage at 2 dpi. Viral positive-strand RNA copies were quantified by RT-qPCR and expressed as genome equivalents (GE) per µg total RNA. The figure was created using Biorender.com.

Oral infection of WT, *Ifnar1^-/-^*, or *Ifnar1^-/-^ Ifnlr1^-/-^* C57BL/6J mice with POWV (10^5^ FFU) resulted in widespread detection of viral RNA in different tissues of mice at 2 dpi (Fig.3B). The highest levels of viral RNA were also present in the spleen following oral infection (Fig.3B). Viral RNA levels in the brain were reduced upon oral infection compared to subcutaneous injection (Fig.3B). Despite being infected with 100 times more viral particles, viral dissemination was less robust across all tissues compared to footpad-injected mice, at least at 2 dpi. No significant difference in viral yield between the two different mouse strains was observed at this timepoint. Type I IFNs restrict rapid dissemination into many peripheral tissues, and more particularly in the spleen. However, the oral route seems less effective in driving systemic dissemination upon disruption of IFN responses than the FP route – despite similar death phenotype (temporal and clinical). This could suggest that the spleen (and the hematopoietic compartment) is a key tissue where disease mechanisms occur when type I IFN signaling is disrupted.

### A CRISPR-KO screen identifies IFI6 as the central effector of the IFN type I and III-driven antiviral responses against TBEV

Since both type I and III IFNs showed antiviral potential against TBOVs *in vitro* (Fig.1) and *in vivo* (Fig.2, 3), we sought to identify the IFN effectors that are involved. We set up a CRISPR/Cas9-KO screening approach targeting 1905 genes that were previously identified as ISGs in a variety of human cell lines and primary immune cells (Fig.4A) (42). To maximize the output of the screen, we performed it both in Caco2 cells treated with IFNλ1 and in DAOY cells treated with IFNα2, since the respective treatments had more potent protective effects (Fig.1I-K). Cells transduced with the sgRNA library were selected by puromycin treatment, then treated with IFN, and finally infected with TBEV at an MOI of 1. Infected cells were sorted using antibodies binding to orthoflaviviral E protein and DNA was extracted from infected and non-infected control cells to identify integrated sgRNA sequences in each cell fraction (Fig.4A). Next Generation Sequencing coupled to MAGeCK-based statistical analyses (43) led to the identification of sgRNAs targeting 19 genes significantly enriched in infected cells, representing potential antiviral factors (Fig.4B). Comparison of antiviral hits in the two models revealed 5 genes that may be essential for both type I and III antiviral signaling against TBEV (Fig.4B). Three of these genes are key members the JAK/STAT pathway and thus well-known broadly-acting IFN-I/III effectors: *STAT1*, *STAT2* and *IRF9* (15,16). *IFI6* and *KEAP1* were the other two antiviral candidates common to the two stimuli. Antiviral hits that were distinct for the two models included the receptors of the respective IFN type used for the two screens, namely *IFNLR1* for the type III/Caco2 screen and *IFNAR1* for the type I/DAOY screen (Fig.4B), further validating our experimental approach. With *IFITM1* and *IFITM3*, ISGs that were previously observed to demonstrate antiviral effects against TBOVs were also detected here (40). Ten of the antiviral hits, such as *TMEM51* and *COMMD3,* have not been associated with antiviral functions yet. We also identified 10 genes that were significantly depleted in infected cells, representing potential proviral ISGs (Fig. 4C). These proviral hits included genes that are broad-spectrum facilitators of viral infection, such as *ISG15* and *USP18*, which negatively regulate the JAK/STAT pathway (44,45). Several genes with unknown functions in the context of viral infection, such as *SLC35A2* and *MARCKS,* were also identified (Fig.4C).

**Fig. 4.**
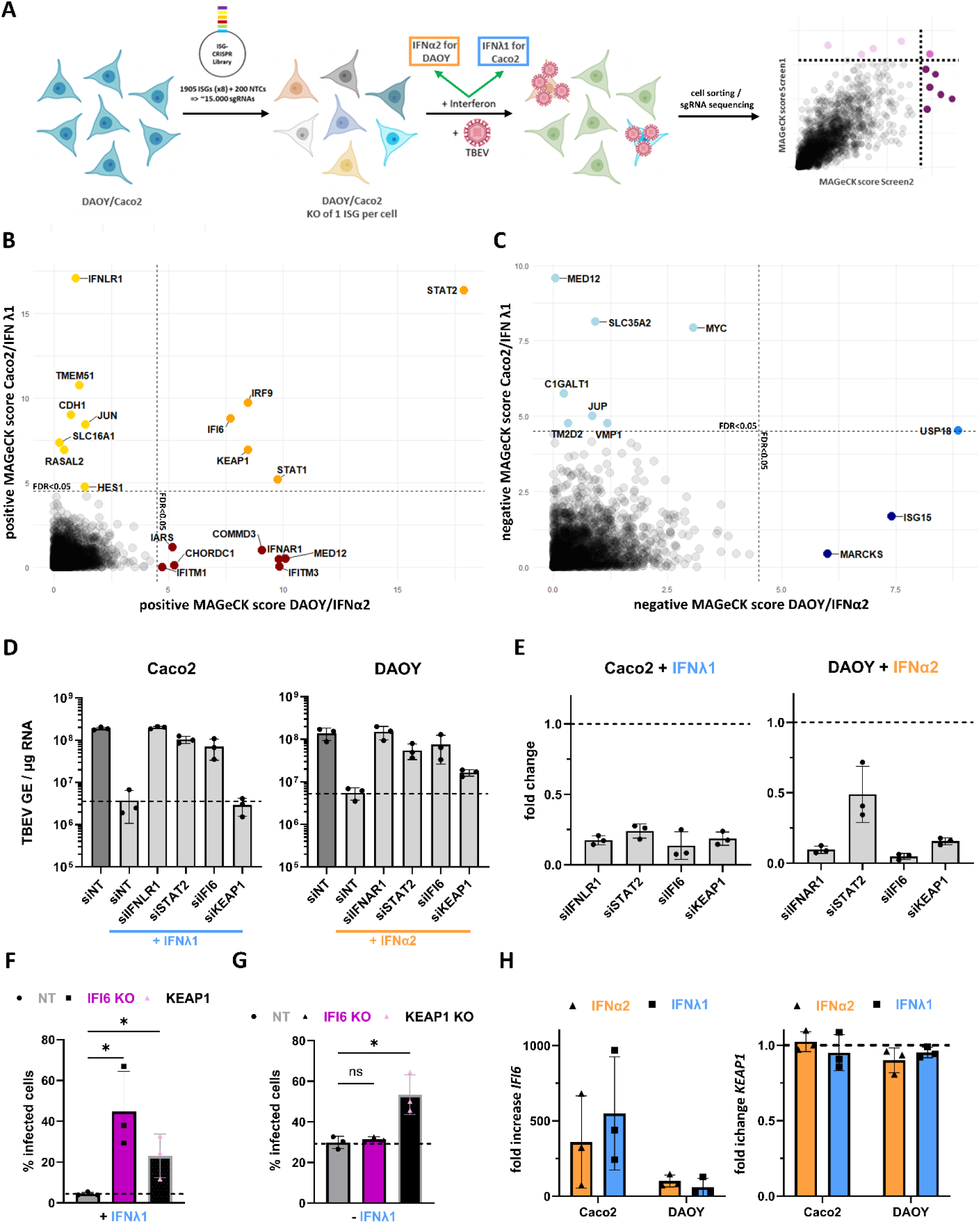
A CRISPR-KO screen identifies IFI6 as the central effector of the IFN type I and III-driven antiviral responses against TBEV. (**A**) Schematic flow chart of the CRISPR-KO screen setup. (**B**) Antiviral and (**C**) proviral hits of the screens as identified through analyses of the resulting NGS data with the MAGeCK package. The cutoff point for screen hits is a FDR of 0.05. (**D**) Caco2 or DAOY cells were issued to siRNA-based knockdown of antiviral screen hits, or transfected with non-targeting siRNA (siNT), treated with IFNλ1 (Caco2, 16h, 20 U/mL) or IFNα2 (DAOY, 16 h, 10 ng/mL), infected with TBEV-Hypr (MOI 1, 24 h) and examined for viral genomic copy numbers *via* RT-qPCR. (**E**) Efficiency of knockdown of expression of target genes in the same IFN-treated samples as in (D) compared to cells transfected with siNT, as assessed by RT-qPCR analysis. (**F**) Caco2 cells with KO of *IFI6* or *KEAP1* or control cells (NT) were treated with IFNλ1 (16 h, 20 ng/mL) and infected with TBEV-Hypr at MOI 0.1 for 24 h. The percentage of infected cells was determined by flow cytometry after staining for orthoflaviviral envelope protein. (**G**) Same setup as (F) without IFN treatment. (**H**) Caco2 or DAOY cells were treated with either IFNα2 (250 U/mL, orange) or IFNλ1 (100 ng/mL, blue) for 16 h and analyzed for mRNA expression levels of *IFI6* and *KEAP1 via* RT-qPCR. Data **(D-H)** are from three independent experiments ± SEM. ns = non-significant, \**P* < 0.05 by unpaired, two-tailed t-test. The figure was created using Biorender.com.

We focused on *IFI6* and *KEAP1,* which were the two antiviral hits that were common to the two screens. To further investigate their antiviral potential, their expression was knocked down using pools of 4 siRNAs. Negative controls were non-targeting siRNAs. Pools of siRNAs targeting *IFNAR1* and *IFNLR1*, which are expected to neutralize IFN-I and IFN-III signaling, respectively, served as positive controls. Pools of siRNAs targeting *STAT2* were also included in the analysis. Following siRNA transfection, DAOY cells were treated with IFNα2 and Caco2 cells with IFNλ1 for 16 hours and then infected with TBEV for another 24 hours. Analysis of viral RNA yield by RT-qPCR, revealed, that, as expected, intracellular viral RNA levels were almost rescued to the level of non-treated cells in the presence of siRNAs targeting IFN receptors and *STAT2* (Fig.4D). RT-qPCR analyses showed that the 4 siRNA pools were reducing the expression of their respective target genes, resulting in a 2-10-fold reduction of mRNA transcripts (Fig.4E). Reduced expression of *IFI6* enhanced TBEV RNA yields in both cell types as compared to IFN-treated cells transfected with control siRNAs (Fig. 4D), confirming that IFI6 significantly contributes to the antiviral state against TBEV, independent of cell- or IFN-type. Replication of TBEV increased in DAOY cells expressing reduced levels of *KEAP1* and pre-treated with IFNα2 (Fig.4D). However, it had no apparent effect on viral replication in IFNλ1-treated Caco2 cells (Fig.4D).

To further investigate the potential effect of *KEAP1* and *IFI6* on TBEV replication, we generated KO cells using lentiviral vectors expressing Cas9 and gRNAs. Caco2 cells expressing non-targeting gRNAs were also produced as control cells. Cells were infected with TBEV and analyzed for the presence of the viral E protein by flow cytometric analysis. As expected from the 2 screens (Fig. 4B) and siRNA-based validation experiments (Fig.4D), significantly more IFI6 KO cells were positive for the viral E protein than control cells upon IFNλ1 treatment (Fig.4F). A significant rescue of viral protein production was also observed in *KEAP1* KO cells (Fig.4F). However, the increase of infection in *KEAP1* KO cells as compared to control cells was independent of prior IFN-treatment (Fig.4G), suggesting that the *KEAP1*-induced antiviral effect is not linked to the IFN-induced antiviral state. Analysis of mRNA levels of *IFI6* and *KEAP1* revealed that *IFI6* was upregulated by IFNα2 and IFNλ1 treatment in both DAOY and Caco2 cells (Fig.4H), thus qualifying as a genuine ISG in these cell types. However, *KEAP1* mRNA abundance remained unchanged upon treatment with IFNα2 or IFNλ1 in both cell types (Fig. 4H), indicating that it is not an ISG in these two cell types.

The screens thus recovered a set of genes with previously described antiviral potential, including genes crucial for IFN-signaling (*IRF9*, *STAT1* and *STAT2*), validating our approach. Furthermore, they identified several poorly characterized host factors potentially modulating TBEV infection in human cells (such as TMEM51 and MARCKS) and IFI6 as a potent effector of both IFN families.

### Diminishing *IFI6* expression impairs the antiviral potential of the IFN type I and III responses against TBEV

Other CRISPR/Cas9 screens previously identified IFI6 as a major player in the type I IFN-dependent antiviral defense against several mosquito-borne orthoflaviviruses, including ZIKV and YFV (46,47). So far, its role in IFN-III signaling has not been explored, and its effect on the replication of TBOVs has not been investigated either. Moreover, the mechanism underlying its antiviral activities is poorly understood.

To confirm the anti-TBEV function of IFI6 in both IFN-I and -III signaling, a pool of four siRNAs was used to assess the effect of reduced *IFI6* expression on viral replication in Caco2 and DAOY cells after treatment with IFNλ1 or IFNα2. Flow cytometry analysis using anti-E antibodies confirmed that IFI6 anti-TBEV activities were dependent on the presence of IFN in both cell lines (Fig. 5A). Similarly, a rescue of the production of infectious virions was observed in cells expressing reduced levels of IFI6, independently of the type of IFN used for pre-treatment and the cell type (Fig. 5B). The siRNA-mediated knockdown reduced the amount of IFI6 mRNA by about 70% in Caco2 cells and by about 90 % in DAOY cells, independently of the type of IFN used for pretreatment (Fig.5C). This validates the used siRNAs as restrictors of *IFI6* expression even in the presence of IFNs.

**Fig. 5.**
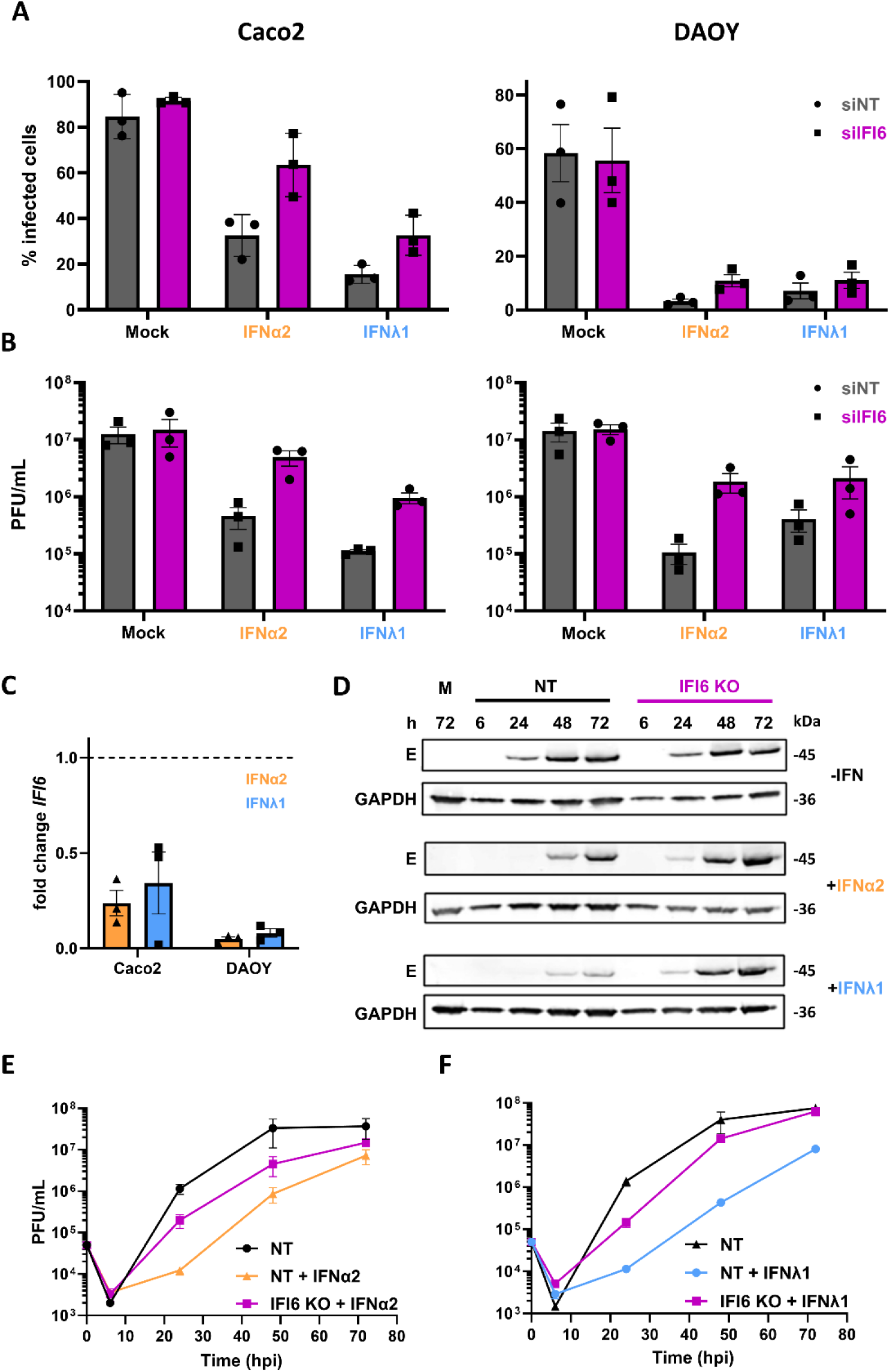
Diminishing IFI6 gene expression impairs the antiviral potential of the IFN type I and III responses against TBEV. **(A)** Cytometry-based analysis of cells stained for orthoflaviviral E protein after siRNA-based knockdown of IFI6 (siIFI6) using a pool of four different siRNAs, or non-targeting control (siNT) and subsequent treatment with either IFNα2 (50 U/mL) or IFNλ1 (20 ng/mL) for 16 h before infection with TBEV-Hypr (MOI 1) for 24 h. **(B)** Same experimental setup as (A) followed by estimating viral titers in the supernatant by plaque assay. **(C)** Reduction of *IFI6* mRNA in Caco2 or DAOY cells transfected with a pool of four siRNAs targeting *IFI6* (siIFI6) expression, or non-targeting siRNAs (siNT) and issued to IFN treatment (IFNα2: 50 U/mL, orange, or IFNλ1: 20 ng/mL, blue) for 16h, as identified by RT-qPCR analysis. **(D)** Western blot analysis of Caco2 cells with stable knockout of IFI6 (IFI6 KO) or control cells transduced with a lentivirus encoding a non-targeting sgRNA (NT) after treatment with either IFNα2 (50 U/mL) or IFNλ1 (20 ng/mL) for 16 h before infection with TBEV-Hypr (MOI 1) for the indicated times. The abundance of viral protein was determined by staining with an antibody against orthoflaviviral E protein. The data is representative of two independent experiments. **(E)** Plaque assay-based titration of supernatants of Caco2 cells with a targeted knockout of IFI6 (IFI6 KO) or control cells (NT) after treatment with IFNα2 (50 U/mL) for 16 h and infection with TBEV-Hypr (MOI 1) for the indicated time. **(F)** Same experimental setup as (F) but treatment with IFNλ1 (20 ng/mL) for 16 h. Data **(A,B,E,F)** are from three independent experiments ± SEM.

The effect of IFI6 on viral replication was further assessed in unstimulated and stimulated IFI6-KO Caco2 cells at different times post-infection. Western blot analysis revealed that the expression of the viral E proteins was increased in cells lacking IFI6, stimulated with IFNλ1 or IFNα2, as compared to control cells that received non-targeting gRNAs (Fig.5D). In agreement with the Western blot analysis, stimulated Caco2 cells lacking IFI6 were producing more infectious viral particles than control cells (Fig.5E-F). The level was almost rescued to the level of control cells, suggesting that IFI6 is an essential contributor to the innate immune response against TBEV. Together, these data confirm that IFI6 plays a central role in the anti-TBEV signaling induced by both type I and III IFNs.

### IFI6 prevents the replication of diverse tick- and mosquito-borne orthoflaviviruses

To further characterize the antiviral activity of IFI6, we produced Caco2 and DAOY cell lines stably expressing a C-terminally 3xFLAG-tagged version of IFI6 (IFI6-FLAG), as well as control Caco2 and DAOY cells stably expressing GFP. The expression of IFI6-FLAG in the generated cell lines was detectable by Western Blot analysis (Fig.6A). *IFI6* mRNA levels exceeded *IFI6* expression in GFP control cells treated with IFN, although less than 1 log in both cell lines with the concentrations of IFN used (Fig. 6B).

**Fig. 6.**
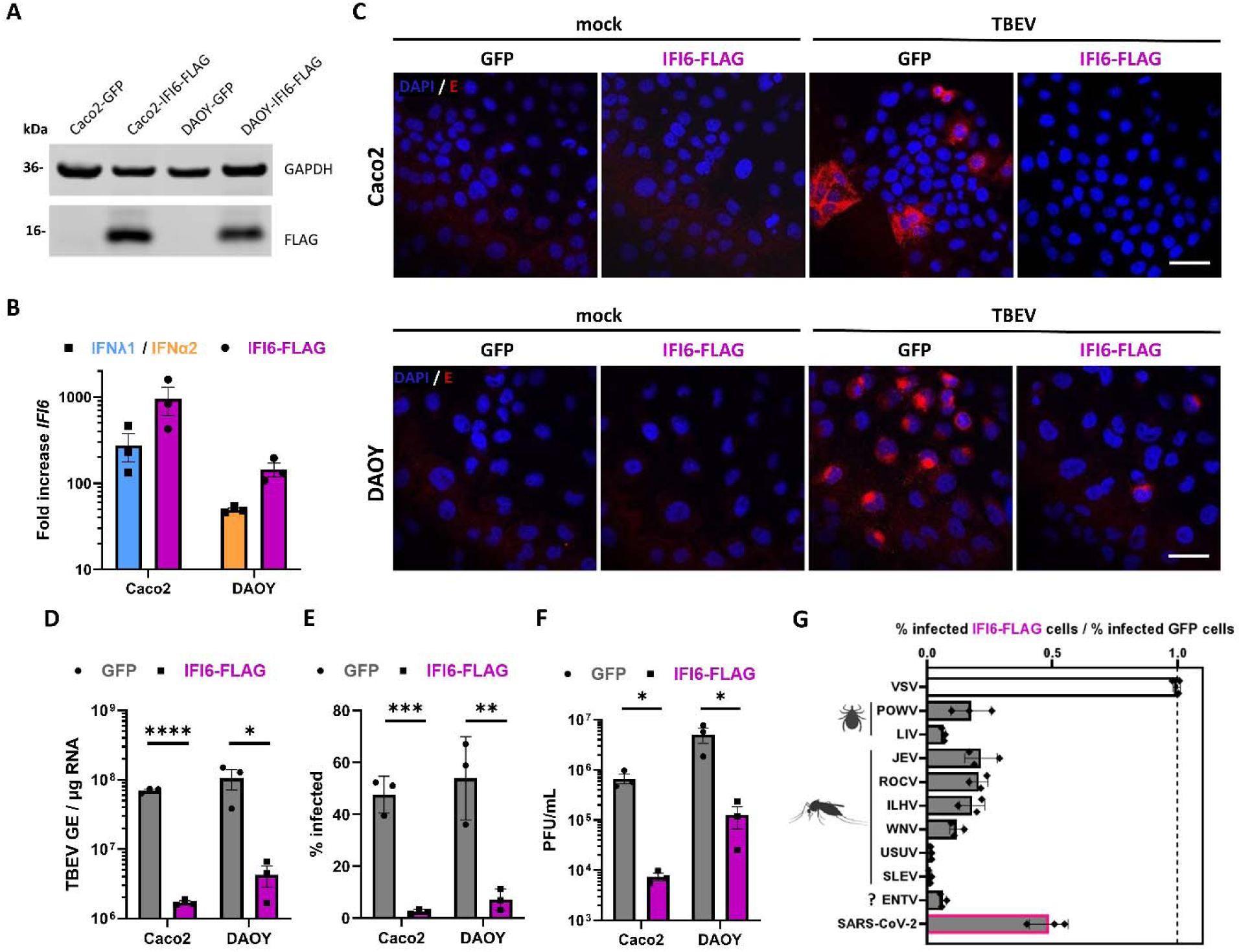
Overexpression of IFI6 prevents replication of diverse tick- and mosquito-borne orthoflaviviruses. **(A)** Western blot analysis of Caco2 or DAOY cells stably expressing IFI6-FLAG or GFP. The abundance of IFI6-FLAG was determined by staining with an antibody against FLAG protein. The data is representative of three independent experiments. **(B)** Expression levels of *IFI6* mRNA in Caco2 or DAOY cells either stably expressing IFI6-FLAG, or after treatment with IFN (Caco2: IFNλ1, 100ng/mL, blue, or DAOY: IFNα2, 250U/mL, orange) for 16h, as identified by RT-qPCR analysis. **(C)** Confocal images of either Caco2 or DAOY cells stably overexpressing GFP or IFI6-FLAG 24 h after mock-infection or infection with TBEV-Hypr (MOI 0.1). Cells were fixed and stained for orthoflaviviral envelope protein (E) to assess their infection status. Images are representative of two independent experiments. Scale bar 30 µm. **(D-F)** Analysis of Caco2 or DAOY cells overexpressing GFP or IFI6-FLAG 24 h after infection with TBEV-Hypr (MOI 0.1) through either RT-qPCR-based estimation of viral genomic copy numbers **(D)**, cytometric assessment of cells positive for stained for orthoflaviviral envelope protein **(E)** or plaque assay-based titration to determine the abundance of infectious virions in the supernatant **(F)**. **(G)** Caco2 cells overexpressing GFP or IFI6-FLAG were infected with either VSV, orthoflaviviruses, or SARS-CoV-2 (MOIs: VSV:0.01; POWV:5; LIV:0.1; JEV:1; ROCV:0.1; ILHV:0.1; WNV:1; USUV:1; SLEV:1; ENTV:1; SARS-CoV-2:0.003) for 24 h and their percentage of infection determined through flow cytometry analysis after specific staining for VSV-G, orthoflaviviral envelope, or SARS-CoV-2 nucleocapsid protein. Transmission vectors are indicated for orthoflaviviruses (ticks, mosquitoes, or unknown (?)). Data **(D-G)** are from three independent experiments ± SEM. ns = non-significant, *P < 0.05, **P < 0.01, ***P < 0.001, and****P < 0.0001 by unpaired, two-tailed t-test.

Confocal imaging revealed that expression of IFI6-FLAG resulted in a reduction of cells positive for viral E proteins 24 h post-TBEV infection in both cell lines, as compared to GFP control cells (Fig.6C). Viral RNA yields were also reduced in DAOY and Caco2 cells expressing IFI6-FLAG (Fig.6D). In agreement with these results, cytometry analysis showed that around 10 times less cells were positive for E proteins in the presence of IFI6-FLAG, as compared to GFP (Fig. 6E). Finally, the release of infectious virions was also significantly reduced in cells expressing IFI6-FLAG (Fig.6F), further validating the antiviral activity of IFI6.

To evaluate the antiviral effect of IFI6-FLAG on other viruses, we conducted infection experiments with an array of diverse orthoflaviviruses transmitted to humans by ticks (POWV and LIV) or mosquitoes (Japanese encephalitis virus (JEV), Rocio virus (ROCV), Ilheus virus (ILHV), West Nile virus (WNV), Usutu virus (USUV), and St. Louis encephalitis virus (SLEV)). We also included Entebbe bat virus (ENTV), an orthoflavivirus isolated from a bat of the species *Tadarida limbata* whose transmission vector has not been identified yet, but that is closely related to YFV (48). Moreover, we tested whether IFI6 was active against Vesicular stomatitis virus (VSV), a negative-sense RNA virus belonging to the *Rhabdoviridae* family that replicates in the cytoplasm (49) as a non-orthoflaviviral control. Flow cytometry analysis using an antibody against the viral protein G revealed that VSV replication was not affected by IFI6 expression (Fig.6G). By contrast, IFI6 significantly reduced the number of E-positive cells in the context of infection with all orthoflaviviruses tested (Fig.6G). Of note, among the orthoflaviviruses tested, USUV and SLEV were the most sensitive to IFI6-overexpression (Fig. 6G). Finally, we assessed the effect of IFI6 overexpression on SARS-CoV-2 replication since two large-scale loss-of-function screens identified it as a potential SARS-CoV-2 restriction factor (50,51). The number of cells positive for the viral protein N was reduced by about 50% when IFI6 was overexpressed (Fig. 6G), indicating that IFI6 antiviral activity is not specific to orthoflaviviruses.

Our experiments thus indicate that IFI6 has the ability to potently restrict the replication of a wide array of orthoflaviviruses, as well as at least one coronavirus, in human cells and acts independently of other ISGs.

### IFI6 is an ER-resident protein that restricts a post-entry step of TBEV replication

To assess the localization of IFI6 within the cell, we tested several commercial antibodies in different assays. None of them gave us satisfactory results. Previous studies have reported a mitochondrial localization of stably overexpressed IFI6-FLAG in human MCF-7 breast cancer cells (52,53), while others described IFI6-3xFLAG as an ER-based protein in human hepatoma Huh7.5 cells in which endogenous IFI6 was replaced by a tagged version using CRISPR knock-in approaches (46). Anti-FLAG staining in DAOY cells stably expressing IFI6-FLAG revealed that the antiviral factor colocalized with the ER marker calreticulin, while no overlap in signal was identified with the mitochondrial marker TOM70 (Fig.7A, B).

**Fig. 7.**
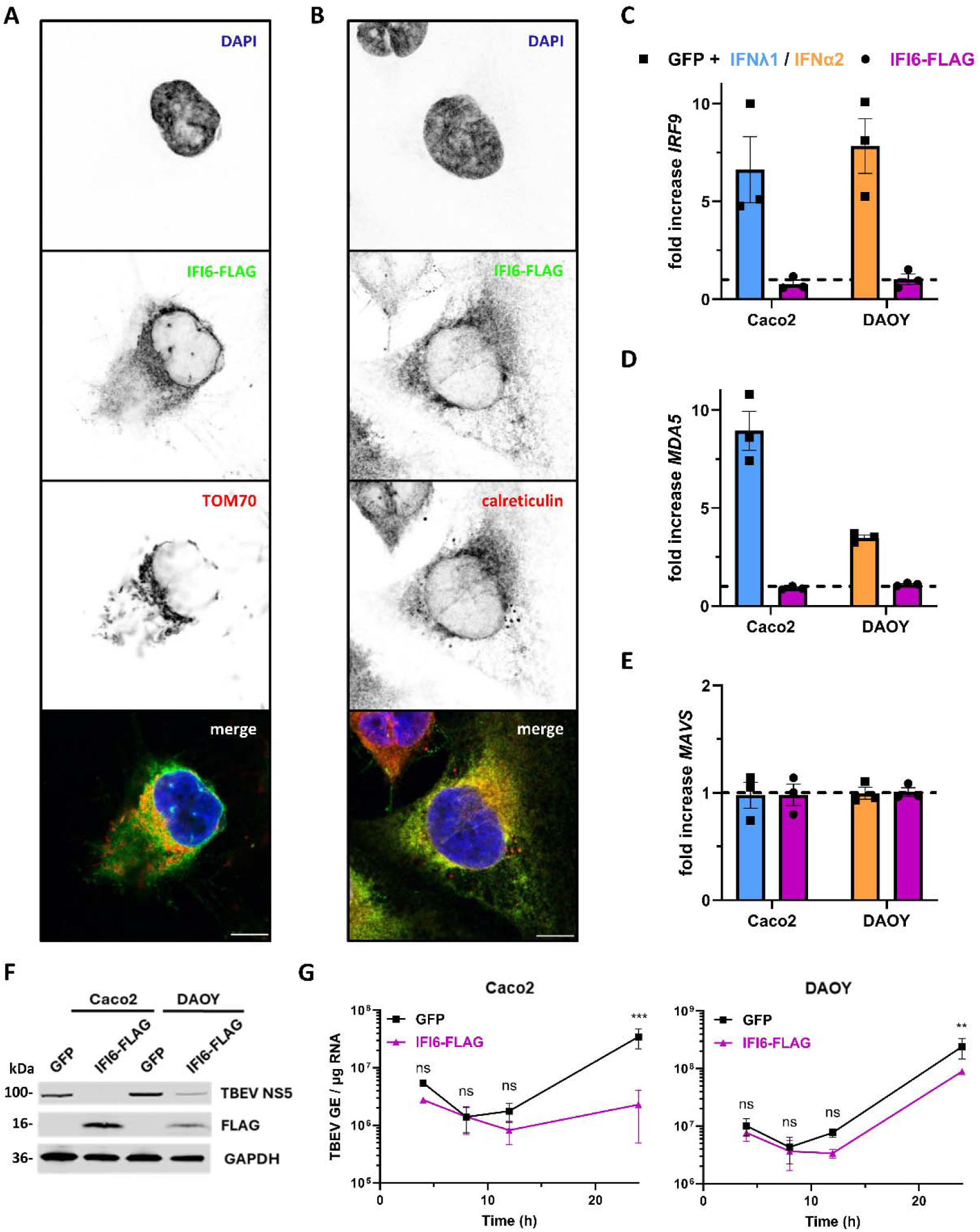
IFI6 is an ER-resident protein that restricts a post-entry step of TBEV replication. (A, **B)** Confocal microscopy analysis of DAOY cells stably overexpressing IFI6-FLAG. Cells were fixed and stained with antibodies for FLAG and mitochondrial marker TOM70 **(A)** or ER-marker calreticulin **(B)**. Scale bar 10µm. **(C-E)** Analysis of *IRF9* (C), *MDA5* (D), or *MAVS* (E) mRNA levels in Caco2 or DAOY cells ectopically expressing GFP or IFI6-FLAG, *via* RT-qPCR. GFP expressing cells were treated with IFN (Caco2: 10 ng/mL IFNλ1, blue; DAOY: 20 U/mL IFNα2, orange) for 16 h. Gene expression was normalized to GAPDH and mRNA levels in mock-treated, GFP-expressing cells. **(F)** Western blot analysis of Caco2 or DAOY cells either ectopically expressing GFP or IFI6-FLAG that were electroporated with *in vitro* synthesized RNA derived from a TBEV Neudoerfl replicon 24 h after electroporation. Whole cell lysates were analyzed by staining with antibodies against the indicated proteins. (n=2). **(G)** Caco2 or DAOY cells, either ectopically expressing GFP or IFI6-FLAG, were electroporated with *in vitro* synthesized RNA derived from a TBEV replicon. At the indicated timepoints, the relative amounts of viral RNA were measured by RT–qPCR analysis of TBEV-NS5 copy numbers per μg of total cellular RNA. Data **(C-E, G)** are from three independent experiments ± SEM. ns = non-significant, \*\**P* < 0.01, and \*\*\**P* < 0.001 two-way ANOVA with Sidak correction.

A recent report suggested an interaction between swine-IFI6 and the NS4A protein of JEV in human embryonic kidney 293T cells, potentially impeding the involvement of NS4A in remodeling of ER-membranes (54). To assess whether TBEV NS4A, or any other viral proteins, could be targeted by IFI6, 293T cells were transfected with FLAG-tagged versions of the 10 individual proteins of TBEV and a C-terminally HA-tagged IFI6. Co-immunoprecipitation assays failed to detect an interaction between viral proteins and IFI6 (Fig.S1A), suggesting that IFI6 does not elicit its antiviral potential *via* an interaction with a viral protein.

Since IFI6 restricts unrelated viruses (Fig. 6G), we assessed whether its activities could be linked to the IFN response. Monitoring the expression levels of three innate immune genes (*IRF9*, *MDA5*, and *MAVS)*, illustrated that while *MAVS* was neither influenced by IFN-treatment, nor IFI6-FLAG expression in Caco2 or DAOY cells, *IRF9* and *MDA5* expression were upregulated by IFNα2 but not influenced by *IFI6* overexpression (Fig.7C-E). These data suggest that IFI6 does not achieve its antiviral potential by modulating the expression of innate immune genes.

To determine whether IFI6 affects early or late steps of TBEV replication, we took advantage of a replicon generated from the NS protein sequences of the Neudoerfl strain (55). DAOY and Caco2 cells stably expressing either IFI6-FLAG or GFP were electroporated with *in vitro* synthesized viral RNA generated from the replicon. In this context, viral replication bypasses early steps (viral attachment, entry, fusion in endosomes, and cytoplasmic uncoating). Analysis of the expression of NS5 in cells transfected with the replicon revealed reduced NS5 levels in DAOY cells and an absence of expression in Caco2 cells after 24 h (Fig.7F). Of note, Caco2 cells expressed more IFI6-FLAG than DAOY cells (Fig.7F), which could explain the more pronounced effect on NS5 expression. RT-qPCR analysis of viral RNA yields performed at different times post-electroporation revealed that viral replication was reduced after around 24 h in both cell lines expressing IFI6-FLAG compared to GFP-expressing control cells (Fig.7G). These data indicate that IFI6 acts at a late step of viral replication.

Together, our findings suggest an antiviral effect of IFI6 by suppressing viral genome replication at a post-entry stage, without interacting with viral proteins.

## DISCUSSION

Our results corroborate the antiviral activity of the type I and III IFN signaling pathways against TBOVs in two human cell lines derived from tissues with physiological relevance to TBOV pathogenesis. A parallel CRISPR/Cas9-KO screen identified IFI6 as a crucial effector of both IFN families in anti-TBEV response. IFI6 likely acts independently of other ISGs and targets a post-entry step shared by various members of the *Orthoflavivirus* genus. In addition to that, IFI6 restricts the replication of at least one member of the *Betacoronavirus* genus.

The importance of type I IFN signaling to counteract TBOV replication in human cells is well established (3,6). In line with our results, the protective effect of pretreatment with type I IFNs was documented for several members of the TBOV family in a variety of cell lines (27,30,31,34). However, to our knowledge, the protective effect of type III IFN pretreatment was only investigated in one study that states that pretreatment of DAOY cells with IFNλ1 does not protect against TBEV replication (30). This is in stark contrast with our results showing the antiviral potential of IFNλ1 against different TBOVs in two human cell lines. Since the experimental setup did not differ in IFN-type nor concentration, the lack of IFNλ1-induced protection from TBEV in DAOY cells might stem from the 5-fold higher MOI previously used (30), potentially overcoming the IFNλ1-induced antiviral state. Several other studies, however, reported antiviral properties of the IFNλ family against diverse mosquito-borne orthoflaviviruses, suggesting that other members of the genus are also sensitive to the antiviral effects of these cytokines (56,47,57–63). The upregulation of type III IFNs following infection with TBOVs was reported in *in vitro* experiments but also in patients infected with TBEV and linked to a milder outcome of disease (30,64,65). Furthermore, studies performed on the Polish and Russian populations suggested that, amongst others, polymorphisms in the *IFNL3/IL28B* gene are associated with a predisposition to severe forms of TBE (64,66). Treatment of patients with IFNλ is currently successfully explored in the context of a variety of viral infections in humans, at present focusing on hepatic and respiratory viruses (67–73). Compared to type I IFNs, previously used for treatment of viral infections, type III IFNs trigger less expression of inflammatory cytokines, like IL-6 and TNFα (21,67,74). IFNλs may thus represent an interesting option to treat patients suffering from neurotropic TBOV infection.

Only the impact of type I IFN signaling on TBOV infection was studied *in vivo* so far (29,33,41,75,76). Furthermore, data on POWV infection in mouse models is scarce. Our data underlines the importance of type I IFN signaling in restricting early viral dissemination and delaying lethal POWV-induced disease. The rapid and fulminant disease we observed in the *Ifnar1^-/-^*and *Ifnar1^-/-^ Ifnlr1^-/-^* mice is unlikely to reflect the typical neurological disease reported for POWV-infected individuals, but rather a systemic ‘cytokine storm’. The additional depletion of IFN-III signaling did not alter the probability of survival for subcutaneously infected C57BL/6J mice. Viral dissemination to spleen and brain as early as day 2 was reported for *Ifnar1^-/-^*C57BL/6J mice intraperitoneally injected with Langat virus (LGTV) (29), complementing our observations for infection of these tissues by POWV after two days. Infection of the liver, although not reported for murine models challenged with POWV, complements reports of TBOV tropism for this tissue in infected patients (77). Importantly, subcutaneous infection with POWV resulted in the detection of viral RNA in the small intestine of mice depleted for IFN-signaling. This is in line with studies showing that TBEV also targets GI tissue in BALB/c or C57BL/6J mice following intravenous or subcutaneous injection, respectively and with GI complications in patients during acute TBOV infection (78,79).

While transmission through ingestion was established for TBEV in a C57BL/6 mouse model and is frequently observed in patients, it remains unclear whether other TBOVs are able to exploit this way of transmission (33,80,81). The only study investigating oral transmission of POWV in animals revealed that infection of a lactating goat with POWV did not result in clinical signs or detectable viremia in the animal but resulted in the development of specific antibodies in its kid, suggesting milk-borne transmission of POWV (8), which is in line with our data. Moreover, vertical transmission of LGTV through breast milk in A129 mice was reported, and also cases of mother-to-child transmission of TBEV *via* breast milk were suggested for humans (36). These findings imply that potentially all TBOVs are transmittable from infected mothers to their children during breastfeeding, which needs to be considered for management of breastfeeding after tick bites in TBOV-endemic areas.

The trend towards earlier lethality in orally infected *Ifnar1^-/-^ Ifnlr1^-/-^* mice compared to mice depleted of only IFN-I signaling supports a greater importance of type III IFN-dependent antiviral signaling at mucosal surfaces than in the skin, as suggested for other viral families (82). However, the observed trend is modest, and no differences in viral titer in the sera of infected *Ifnar1^-/-^* and *Ifnar1^-/-^ Ifnlr1^-/-^*mice were observed at 2 dpi. This fits the comparable loads of viral RNA extracted from diverse tissues in orally infected mice and suggests a minor role of type III IFN signaling for the antiviral defense system *in vivo*, independently of the infection route. Generally, infection with POWV through oral gavage resulted in significantly lower systemic viral titers and abundance of POWV RNA in the examined tissues compared to footpad-injected mice. These results suggest more efficient viral dissemination routes following subcutaneous infection than oral infection of POWV. However, when C57BL/6J *Ifnar1^-/-^*mice were subcutaneously infected with LGTV (10^2^ FFU), they succumbed significantly earlier to the virus compared to perorally infected mice (10^4^ FFU) (33). Interestingly, *Ifnar1^-/-^*mice infected with up to 10^6^ FFU of LGTV through oral gavage did not die, suggesting that peroral infection with TBOVs causes more efficient dissemination of the virus than administration *via* oral gavage (33). Similar observations were made for TBEV in WT and *Ifnar1^-/-^* mice (33).

The antiviral state induced by IFNs depends on the activation of a plethora of different ISGs, many of which remain uncharacterized (18). While several approaches successfully identified IFN-I effectors against mosquito-borne orthoflaviviruses (46,47,83–85), effectors against TBOVs are poorly described. Our parallel CRISPR/Cas9-KO screening approach recovered factors previously identified as antiviral effectors against TBOVs, such as *IFITM1* and *IFITM3* (40). Moreover, with *IARS*, *MED12*, *CHORDC1,* and *COMMD3* for the type I screen and *TMEM51*, *CDH1*, *JUN*, *SLC16A1*, *RASAL2,* and *HES1* for the type III screen, we identified several novel genes with potential involvement in the IFN-induced cellular defense mechanism against TBOVs. Further investigations would be required to validate the antiviral activities of these candidates and decipher their mechanisms of action. Of note, we did not identify *RSAD2* (viperin) nor *TRIM5*α, two ISGs previously were reported to counteract TBOV infection in human A549 and/or HEK293T cells, although they were included in the sgRNA library we screened (42,86–89). Their antiviral function may be cell-type specific. The screens further suggested a proviral effect of *USP18*, *ISG15,* and *VMP1*, which agrees with previous experiments performed in human cells infected with other members of the *Flaviviridae* family (44,45). *KEAP1* emerged as a potential IFN-I and IFN-III effector against TBEV. Although further investigations revealed that its action is IFN-independent in Caco2 and DAOY cells, it may be induced by IFNs in other cell types.

Our screens identified *IFI6* as an IFN-I and IFN-III effector for the anti-TBEV response in human cells. Reducing IFI6 expression in the presence of IFN-I/-III enhanced TBEV replication to a level comparable to the inhibition of IFNAR1 or STAT2, suggesting that it largely contributes to the innate immune response against TBEV. We showed that IFI6 restricts the replication of orthoflaviviruses transmitted by ticks, *Culex* and *Aedes* mosquitoes. In line with previous reports (46), our data support the hypothesis of IFI6 being an ER resident and suppressing the replication of orthoflaviviruses at a post-entry step. We observed an anti-SARS-CoV-2 effect of IFI6 expression in Caco2 cells, which was unexpected since the replication of OC43, a related betacoronavirus, was unaffected by IFI6 overexpression in Huh7.5 cells (49). These effect on SARS-CoV-2 replication thus challenges the idea of IFI6 solely preventing the formation of orthoflavivirus-specific alterations in the ER membrane (46). IFI6 may thus restrict the replication of unrelated viruses that replicate in the ER, preventing the formation of virally-induced ER remodeling (46), a process that is key for orthoflaviviral and betacoronaviral replication (90–92). This re-arrangement of ER membrane structures is mostly driven by several viral proteins, even though the detailed mechanisms of their involvement are yet to be defined (93). How the restructuring of the ER membranes by viral proteins might be influenced by IFI6, however, remains unclear. A direct interaction between the 2K protein of JEV, which lies in the last transmembrane domain of NS4A, and swine IFI6 (sIFI6) was recently suggested (54). Cleavage of the 2K protein from NS4A through the viral NS2B-NS3 protease modulate its potential to induce membrane rearrangements (94,95). This is further illustrated by a mutation in the genome of WNV that prevents cleavage of NS4A-2K completely abrogating viral replication (96). IFI6 may thus block the 2K cleavage and prevent the ER-remodeling potential of the NS4A protein of orthoflaviviruses. However, we did not detect interaction between human IFI6 and TBEV NS4A-2K, or any other viral protein. Furthermore, IFI6 was not able to prevent NS4A-2K cleavage, or any other polyprotein processing events induced by the viral protease (46,47). Together, these results challenge the idea of IFI6 exercising its antiviral potential by inhibiting viral protease activity and subsequent prevention of ER remodelling. This makes further investigation of the mechanisms underlying IFI6 antiviral activities indispensable. Importantly, this is hindered by the fact that no murine ortholog of IFI6 exists, greatly complicating its investigation in an *in vivo* context.

## MATERIALS AND METHODS

### Mice

All animal experiments described in this study were performed in accordance with protocols that were reviewed and approved by the Institutional Animal Care and Use Committee of Boston University (PROTO202100000025 (Breeding protocol) and PROTO202200000070 (BSL3 protocol)). All mice were maintained in facilities accredited by the Association for the Assessment and Accreditation of Laboratory Animal Care (AAALAC). All replication-competent Powassan Virus experiments were performed in a biosafety level 3 laboratory (BSL-3) at the Boston University National Emerging Infectious Diseases Laboratories (NEIDL).

C57BL/6J (Cat. # 000664) and B6(Cg)-Ifnar1tm1.2Ees/J (*Ifnar1^-/-^*mice; Cat. # 028288) mice that are deficient for type I innate immune signaling were acquired from Jackson Laboratories. *Ifnar1^-/-^ Ifnlr1^-/-^* mice that are deficient in type I and III innate immune signaling were kindly provided by Sergei Kotenko at Rutgers University. All mice were maintained at the Animal Science Center at Boston University prior to being moved to the NEIDL for experiments.

Mice in the NEIDL BSL-3 facility were group-housed by sex in Tecniplast green line individually ventilated cages (Tecniplast). Mice were maintained on a 12:12 light cycle at 30– 70% humidity and provided standard water and standard chow diets (LabDiet).

### Cell lines and viruses

Human colorectal adenocarcinoma Caco2-TC7 (American Type Culture Collection (ATCC), HTB-37), human medulloblastoma DAOY (ATCC, HTB-186), Human embryonic kidney (HEK) 293T cells (ATCC, CRL□3216), human glioblastoma U87-MG (ATCC, HTB-14), African green monkey kidney epithelial Vero cells (ATCC), and human hepatoma-derived HuH7 cells (HuH7.5, ATCC) were maintained in Dulbecco’s modified Eagle’s medium (DMEM) (Gibco) containing GlutaMAX I and sodium pyruvate (Invitrogen) supplemented with 10% heat inactivated fetal bovine serum (FBS) (Dutscher) and 1% penicillin and streptomycin (10,000 IU/ml; Thermo Fisher Scientific) at 37°C and 5% CO2.

Experiments with TBEV, POWV, LIV, ZIKV, JEV, ROCV, ILHV, WNV, USUV, SLEV, ENTV, and SARS-CoV-2 were performed in a BSL□3 laboratory, following safety and security protocols approved by the risk prevention service of the Institut Pasteur, Paris, or the NEIDL at Boston University. Experiments with VSV and lentiviruses were performed in a BSL-2+ setting following biosafety regulations of the Institut Pasteur, Paris. VSV Indiana was kindly provided by N. Escriou (Institut Pasteur, Paris). The TBEV-Hypr (001v-EVA134), POWV-LB (001v-EVA124), ROCV (001v-EVA126), ILHV (001v-EVA106), and ENTV (001v-EVA84) were obtained from the European Virus Archive (EVAg; https://www.european□virus□archive.com/). Louping Ill virus (strain LI 3/1; APHA reference Arb126) was a kind gift from Nick Johnson (Animal and Plant Health Agency, Addlestone, Surrey, UK). JEV (strain RP9), SLEV (strain MSI-7) and WNV (strain IS-98-STI) were provided by the biological resource center of the Institut Pasteur. USUV (strain D18-03348) was a kind gift from Nolwenn Dheilly (Institut Pasteur, Paris, France). SARS-Cov-2 (strain BetaCoV/France/IDF0372/2020) was supplied by the French National Reference Centre for Respiratory Viruses hosted by Institut Pasteur (Paris, France). Vero cells were used to produce all viral stocks, and to assess the number of infectious virions produced through plaque assay, as previously described (97). Only POWV-LB virus used for infection of mice was titrated using a focus forming assay on Vero cells. Briefly, Vero cell monolayers in a 24-well plate were infected with serial dilutions of virus stock for 2 h at 37°C, then the supernatant was replaced with a 1:1 mixture of 2xDMEM and CMC and incubated for 6 days at 37°C under a 5% CO2 atmosphere. The overlay was then removed, cells washed with PBS, fixed with 4% PFA at RT for 15 min and then permeabilized and blocked with PBS (1%BSA, 1% Triton-X) at RT for 1 h. Cells were then incubated with PBS (1% BSA) with pan-orthoflaviviral E protein 4G2 antibody (1:2000, Thermo Fisher, NBP52666549) at RT for 1 h while shaking. Monolayers were then washed 3x with PBS and incubated with PBS (1% BSA) holding Li-Cor IRDye® 800CW anti-rabbit antibody (1:5000, Fisher, NC9523609) for 45 min. Cells were washed 3x with PBS and plates imaged using an Odyssey DLx machine (Li-Cor). Viral titers were calculated according to the dilution and plated volume.

### Antibodies and cytokines

The following primary antibodies were used: commercially available antibodies against β□actin (produced in mouse, clone AC□74), FLAG (mouse, M2 F3165□1MG), 4G2 pan orthoflavivirus (mouse, 1:1000, kind gift from Philippe Desprès), anti-TBEV NS5 (34), TOM70 (rabbit, ab289977), calreticulin (rabbit, AB2907), and HA (rabbit, H6908) were obtained and used for immunoblotting at the indicated dilutions. Furthermore, FLAG M2 was used at 1:2000, 4G2 at 1:2000, and anti-NS5 at 1:1000 dilutions for immunofluorescence staining for microscopy analysis. 4G2 pan□orthoflavivirus antibody was used at 1:1000, anti-VSV-G antibody (kind gift from Gert Zimmer) was used at 1:2000, and anti-SARS-Cov-2-NCP-1 antibody (kind gift from Olivier Schwartz (98)) was used at 1:200 dilutions in flow cytometry, respectively.

For secondary staining, the following commercially available antibodies were employed: Alexa Fluor 488 goat anti□mouse IgG (H+L, A11001), Alexa Fluor 647 donkey anti□mouse IgG (H+L, A31571), Alexa Fluor 647 goat anti□rabbit IgG (H+L, A21244); Alexa Fluor 680 goat anti□mouse IgG (H+L, A21058); goat anti□chicken IgY (H+L) Dylight 800 (SA5□10076) and goat anti□rabbit IgG (H+L) Dylight 800 (SA5□35571). All secondary antibodies were used at 1:1,000 dilution for cytometry and immunofluorescence assays and diluted at 1:10,000 for Western blot analysis.

Human recombinant IFNα2 (PBL Biosciences) and IFNλ1 (IL-29, Invitrogen) were used to stimulate cells at the concentrations (diluted in DMEM 10% FBS; 1% P/S) and for the time indicated in the respective figure legends.

### Viral infections

Cells were infected at the MOIs indicated in the respective figure legends by a 2 h incubation in DMEM medium containing 2% FBS at 37°C under a 5% CO2 atmosphere. Cells were then washed with PBS (Gibco), infection medium was replaced by DMEM (10% FBS; 1% P/S) and cells were subsequently incubated at 37°C under a 5% CO2 atmosphere until collected for analysis at the indicated timepoint.

### Transfections

293T cells were transfected using the Trans IT□293 (Mirus) reagent following the protocol provided by the manufacturer. For co immunoprecipitation experiments, 5×10^5^ cells in 12□well plates were transfected with 250□ng of each plasmid, as indicated in the figure.

The ON-TARGETplus siRNAs used for either screen validation or IFI6 knockdown experiments in DAYO and Caco2 cells (Table S1) were purchased from Horizon. Caco2 and DAOY cells were seeded on a 12-well plate (8×10^4^/well) and transfected with Lipofectamine RNAiMAX transfection reagent (Thermo Fisher Scientific) according to the manufacturer’s instructions.

### Immunoblot and immunoprecipitation analyses

Cells were lysed using radioimmunoprecipitation assay (RIPA) buffer (Sigma□Aldrich), supplemented with a protease and phosphatase inhibitor cocktail (Roche). Subsequently, samples were denatured in 4x protein sample loading buffer (Li□Cor Bioscience) under reducing conditions (NuPAGE reducing agent, Thermo Fisher Scientific) when non-infected, or not when infected with orthoflaviviruses. Only 293T cells expressing TBEV NS2A were lysed using the Mem□PER plus membrane protein extraction kit (Thermo Fisher Scientific) and sonicated for 20□minutes at 100% amplitude and 2/2 pulse. Proteins were then separated by SDS–PAGE (NuPAGE 4-12% Bis□Tris gel, Invitrogen) and transferred to nitrocellulose membranes (Bio□Rad) utilizing a Trans□Blot Turbo Transfer system (Bio□Rad). Membranes were blocked with PBS 0.1% Tween 20 (PBS□T) containing 5% milk. Following blocking, membranes were incubated for 1 h at RT or overnight at 4°C with primary antibodies diluted in blocking buffer. Subsequently, membranes were washed 3x with PBS-T and incubated for 45□minutes at room temperature with secondary antibodies (anti□rabbit/mouse IgG (H+L) DyLight 800/680) diluted in blocking buffer, followed by 3x washing of the membranes with PBS-T. Images were acquired using an Odyssey CLx infrared imaging system (Li□Cor Bioscience).

For co□immunoprecipitation analysis, a portion of the cell lysates was incubated overnight with magnetic beads coupled with anti FLAG M2 (Sigma□Aldrich, M8823). Following incubation, immunoprecipitates were washed four times with washing buffer and analyzed by immunoblot as described above.

### RNA extraction and RT-qPCR analyses

Total RNA was extracted from cell lysates utilizing the NucleoSpin RNA II kit (Macherey□Nagel) or the NucleoMag RNA kit (Macherey-Nagel) according to the manufacturer’s instructions and subsequently eluted in nuclease free water. First□strand cDNA synthesis involved 1□μg of total RNA using the RevertAid H Minus Moloney murine leukemia virus (M□MuLV) reverse transcriptase (Thermo Fisher Scientific) with random primers p(dN)6 (Roche). Quantitative real□time PCR was conducted on a Quant Studio 6 Flex real-time PCR system (Applied Biosystems) using SYBR green PCR master mix (Life Technologies). Data were analyzed employing the ΔΔCT method, with GAPDH (glyceraldehyde□3□phosphate dehydrogenase) serving as the normalization reference for human samples. Technical triplicates were performed for each experiment. RT–qPCR primer sequences are detailed in Table S2. Quantification of TBEV and POWV genomes was achieved by extrapolation from a standard curve derived from serial dilutions of plasmids encoding TBEV NS5 (pCi□Neo□NS5 TBEV) or POWV NS5 (pCi-Neo-NS5 POWV).

### Immunofluorescence assays

Cells were fixed with 4% paraformaldehyde (PFA) (Sigma□Aldrich) at RT for 30□minutes, followed by permeabilization with PBS containing 0.5% Triton-X at RT for 15□minutes. Subsequently, cells were blocked for 30□minutes with PBS containing 0.05% Tween and 5% BSA, before being exposed to the specified primary antibodies diluted in PBS containing 0.05% Tween and 2% BSA for 1□h. Following antibody incubation, cells were washed 3x with PBS containing 0.05% Tween. Secondary Alexa Fluor 488 or 647-conjugated antibodies diluted in PBS containing 0.05% Tween and 2% BSA were then applied for 1□h. After secondary antibody incubation, cells were washed 3x with PBS containing 0.05% and once with PBS alone. Nuclei were stained for 15□minutes using PBS/NucBlue (Life Technologies). Following staining, slides were mounted with Prolong gold imaging medium (Life Technologies). Images were captured utilizing a Leica SP8 confocal microscope.

### Flow cytometry

Infected cells underwent fixation using the Cytofix/Cytoperm fixation and permeabilization kit (BD Pharmingen) for 20 min, followed by three washes with the corresponding wash buffer. Subsequently, cells were stained with the anti□E mAb 4G2 primary antibody, diluted in wash buffer, at 4°C for 1□h. After another three washes with wash buffer, cells were stained with secondary Alexa 488 or Alexa 647 antibody for 45□minutes in the dark at 4°C. Data acquisition was performed using the Attune NxT Acoustic Focusing Cytometer (Life Technologies), and analysis was conducted using the FlowJo v10.8.1 software.

### Generation of ISG-KO library cells

Twenty 6-well plates were seeded with 4×106 HEK293T cells in DMEM (10% FBS, 1% P/S). After 24 h, cells were simultaneously transfected with the 667 ng lentiCRISPRv2 ISG library plasmids (https://www.addgene.org/pooled-library/lenticrisprv2-isg-library/), 500 ng plasmids coding for HIV Gag/Pol (psPAX2) and 333 ng of plasmids encoding for the VSVg envelope (pVSV-G opt) per well using Trans IT□293 (Mirus) transfection reagent following the manufacturer’s instructions. Transfection medium was replaced after 24 h, and supernatants were harvested 24 and 48 h later, filtered, concentrated by ultracentrifugation (22,000□g, 4□°C for 1□h), and pooled. To generate the ISG KO library cells, 3×10^7^ Caco2 or DAOY cells were seeded in 12-well plates 24□h before transduction. Each well was then transduced at an MOI of 1 as identified by colony formation titering assays for lentiviruses on Caco2 or DAOY cells following previously described protocols (99). Briefly, concentrated lentivector was diluted in serum-free DMEM, supplemented with 20□µg/mL of DEAE dextran (Sigma, D9885), and cells were spin-infected for 30 min at 1100xg. After 48□h, transduced cells were selected by puromycin treatment for 10 days (Caco2: 10 µg/mL, DAOY: 4□µg/mL; Sigma, P8833).

### CRISPR/Cas9 screens

In total, 1.3□×□10^7^ Caco2 or DAOY cells were treated with IFNλ1 (Caco2, 100 ng/mL) or IFNα2 (DAOY, 100□U/mL). After 16□h, cells were infected with TBEV-Hypr at an MOI of 1 in DMEM (2% FBS, 1% P/S). After 2h, the viral inoculum was removed, washed once with PBS, and cells were maintained in DMEM containing 10% FBS and 1% P/S for 24 h. Thereafter, cells were collected and fixed using the fixation buffer supplied with the Cytofix/Cytoperm™ fixation and permeabilization kit (BD Pharmigen). Fixed cells were washed in cold Cytoperm buffer and then incubated for 1 h at 4□°C under rotation with primary antibody (4G2, mouse, 1:1000) in the same buffer. Incubation with the secondary antibody (Alexa 488 anti-mouse, 1:10000) was performed for 45□min at 4□°C under rotation. Stained cells were resuspended in cold sorting buffer containing PBS, 2% FBS, 25□mM Hepes, and 5□mM EDTA. Cells were sorted into infected and non-infected cells using a BD FACS Aria Fusion machine. Sorted and control (non-infected, not IFN-treated) cells were then centrifuged for 20□min at 2000xg and resuspended in lysis buffer (NaCI 300□mM, SDS 0.1%, EDTA 10□mM, EGTA 20□mM, Tris 10□mM) supplemented with 1% Proteinase K (Qiagen) and 1% RNAse A/T1 (Sigma) and incubated overnight at 65□°C. DNA was isolated with two consecutive phenol-chloroform extractions and recovered by ethanol precipitation. Next, a PCR reaction using barcoded primers to allow for demultiplexing of the samples after sequencing was performed using the Herculase II Fusion DNA Polymerase (Agilent) and DNA oligos as described in (2). The resultant PCR products were purified using a QIAquick PCR Purification kit (Qiagen, 28104) and subjected to a second PCR following primer designs described in (99) before being purified using Agencourt AMPure XP Beads (Beckman Coulter Life Sciences). Next-generation sequencing was performed utilizing the NextSeq 500/550 High Output Kit v2.5 with 75 cycles (Illumina).

### Screen analysis

Reads were demultiplexed using quality-aware fastq demultiplexer v0.1.1. Sequencing adapters were removed using trimmomatic v0.39. The reference library was built using bowtie2 v2.3.5.1. Read mapping was performed with bowtie2 and samtools v2.0.4. Mapping analysis and gene selection were performed using MAGeCK v0.5.9, normalizing the data with default parameters. Data from MAGeCK analyses of the screens are available upon request to the authors.

### Generation of cells with stable KO of IFI6 or KEAP1

Stable KO of IFI6 or KEAP1 in Caco2 or DAOY cells was achieved through transduction with lentivectors harboring sgRNA sequences targeting the specific genes (Table S3) introduced into the lentiCRISPRv2 backbone. The sgRNAs were produced by hybridization of oligonucleotides encoding for the respective sgRNA sequence flanked by sequences resembling restriction sites for the BsmBI enzyme. LentiCRISPRv2 plasmids and the hybridized oligonucleotides were digested with the restriction enzyme and gel purified. The components obtained were then ligated using a T4 ligase for 2 h at RT. The resultant plasmids were transformed into Stbl3 bacteria for amplification and sequenced to ensure the successful incorporation of the sgRNA sequence. After, HEK293T cells plated in a 6-well plate and grown to 70% confluency were simultaneously transfected with the 667 ng PIKA sgRNA library plasmids, 500 ng plasmids coding for HIV Gag/Pol (psPAX2) and 333 ng of plasmids encoding for the VSVg envelope (pVSV-G opt) per well using Trans IT□293 (Mirus) transfection reagent following the manufacturer’s instructions. Transfection medium was replaced after 24 h, and supernatants were harvested 24 and 48 h later, filtered, and pooled. To generate the KO cells, a 6-well plate with Caco2 or DAOY cells was incubated with lentivector-holding supernatant diluted 1:2 in serum-free DMEM for 2 h before replacing the inoculum with DMEM (10% FBS, 1% P/S). After 48 h, transduced cells were selected by puromycin treatment for 10 days (Caco2: 8 µg/mL, DAOY: 4□µg/mL; Sigma, P8833).

### Generation of IFI6-Flag expression plasmid and lentiviral vectors

IFI6-Flag expression plasmid was generated by cloning the IFI6 coding sequence from IFN-stimulated DAOY cells’ cDNA into p3XFLAG-CMV™-14 vector (Sigma) using EcoRI/KpnI restriction sites (Fwd: 5’-TCTAGTGAATTCCCACCATGCGGCAGA AGGCGGTATCGC -3’, Rev: 5’-GACTGGTACCTCCTCATCCTCCTCACTATC-3’).

The generated IFI6-3xFLAG plasmid was used to transfer the IFI6-3xFLAG cassette into the pFlap-Ubc-nLuc-IRES-Puro lentiviral vector (kind gift of Pierre Charneau). PCR primers were designed to add BclI/XhoI restriction sites to IFI6-3XFLAG coding sequence (Fwd: 5’-CTCTAGAGGATCCCGGGCTG-3’, Rev: 5’-GAGAGGCTCGAGTCTCACTACTTGTCATCGTCA-3’), and the amplicon was cloned into pFlap-Ubc-IRES-Puro lentiviral vector digested with BamHI/XhoI to generate the pFlap-Ubc-IFI6-3xFLAG-IRES-Puro plasmid. Amplicon was cloned in pFlap-Ubc-IRES-Puro using BamHI/XhoI restriction sites.

The HA tagged version of IFI6 was obtained amplifying IFI6 from the IFI6-3XFLAG plasmid, with primers designed to add BclI/XhoI restriction sites along with a C-terminal HA tag (Fwd: 5’-TCTAGTTGATCAGCCACCATGCGGCAGAAGGCGGT-3’, Rev: 5’-GAGAG GCTCGAGTCACTAAGCGTAATCTGGAACATCGTATGGGTAAGCCCGGGATCCTCTAGAGTC-3‘) and amplicon was cloned into pFlap-Ubc-IRES-Puro lentiviral vector digested with BamHI/XhoI to generate the pFlap-Ubc-IFI6-HA-IRES-Puro plasmid used for transfection.

Control GFP expressing lentiviral vector was obtained using the same cloning strategy, amplifying GFP from pEGFP-C1 plasmid (Clontech) (Fwd: 5’-TCTAGTTGATCAGCCACCATGGTGAGCAAGGGCGA-3’, Rev: 5’-GAGAGGCTCGAGTCTTACTTGTACAGCTCGTCCAT-3’). All plasmids were propagated in Stbl3 *E. coli* cells and verified by sequencing (Eurofins Genomics).

### Generation of stable cell lines

HEK293T cells plated in a 6-well plate and grown to 70% confluency were simultaneously transfected with 1.6 µg of either p6.16-Lenti-GW-TMEM51, pFlap-Ubc-IFI6-3xFLAG-IRES-Puro pFlap-Ubc-IFI6-HA-IRES-Puro or pFlap-Ubc-EGFP-IRES-Puro, 1 µg plasmids coding for HIV Gag/Pol (psPAX2) and 400 ng of plasmids encoding for the VSVg envelope (pVSV-G opt) per well using Trans IT□293 (Mirus) transfection reagent following the manufacturer’s instructions. Transfection medium was replaced after 24 h, and supernatants were harvested 24 and 48 h later, filtered, and pooled. To generate the cells stably expressing IFI6-FLAG, a 6-well plate with Caco2 or DAOY cells was incubated with lentivector-containing supernatant diluted 1:2 in serum-free DMEM for 2 h before replacing the inoculum with DMEM (10% FBS, 1% P/S). After 48□h, transduced cells were selected by zeocin treatment for TMEM51-expressing cells (Caco2: 300 µg/mL, DAOY: 200□µg/mL, Sigma) or puromycin treatment for IFI6-3xFLAG or IFI6-HA and GFP expressing cells (Caco2: 10 µg/mL, DAOY: 4□µg/mL, Sigma) for 10 days.

HEK293T cells plated in a 6-well plate and grown to 70% confluency were simultaneously transfected with 1.6 µg of either p6.16-Lenti-GW-TMEM51, pFlap-Ubc-IFI6-3xFLAG-IRES-Puro or pFlap-Ubc-EGFP-IRES-Puro, 1 µg plasmids coding for HIV Gag/Pol (psPAX2) and 400 ng of plasmids encoding for the VSVg envelope (pVSV-G opt) per well using Trans IT□293 (Mirus) transfection reagent following the manufacturer’s instructions. Transfection medium was replaced after 24 h, and supernatants were harvested 24 and 48 h later, filtered, and pooled. To generate the KO cells, a 6-well plate with Caco2 or DAOY cells was incubated with lentivector-containing supernatant diluted 1:2 in serum-free DMEM for 2 h before replacing the inoculum with DMEM (10% FBS, 1% P/S). After 48□h, transduced cells were selected by zeocin treatment for TMEM51-expressing cells (Caco2: 300 µg/mL, DAOY: 200□µg/mL, Sigma) or puromycin treatment for IFI6-3xFLAG and GFP expressing cells (Caco2: 10 µg/mL, DAOY: 4□µg/mL, Sigma) for 10 days.

### *In vitro* transcription and electroporation of TBEV replicons

TBEV pTND/ΔME replicon plasmids (kindly provided by Franz X. Heinz *via* Karin Stiasny) (55) were linearized *via* NheI digestion and blunt ended using the Quick Blunting Kit (New England Biolab). 5 µg of purified DNA template were utilized for T7-mediated *in vitro* transcription employing the RiboMAX large□scale RNA production system T7 (Promega), with the addition of 40 mM cap analog (Ribo m7G Cap, Promega) as proposed by the manufacturer’s protocol. After treatment with RQ1 DNase (Promega), RNA was purified using an RNA clean-up kit (Macherey□Nagel). *In vitro* synthesized TBEV replicon RNA was then delivered into Caco2 or DAOY cells *via* electroporation. Here, 2×106 cells were trypsinized, washed 3x in cold PBS, resuspended in 200 µL cold PBS, and electroporated with 5□μg of RNA in 0.4-cm electroporation cuvettes (Biorad), using a 950□μF and 260□V pulse with the Genepulser system (Biorad). Following electroporation, cells were collected in 3.2□ml of warm medium, split into four wells of a 12-well plate, and incubated at 37°C under standard conditions.

### Mouse infection and monitoring

C57BL/6J, *Ifnar1^-/-^*, and *Ifnar1^-/-^ Ifnlr1^-/-^* adult male and female adult mice (6-30 weeks of age) were infected through subcutaneous injection in the foodpad with POWV-LB (10^3^ FFU) diluted in 50 µl of PBS or through oral gavage with POWV-LB (10^5^ FFU). Clinical manifestations of disease were monitored daily, and signs of clinical disease progression were recorded through weighing, clinical scoring, and temperature measurements using a UID temperature implantable probe (Unified Information Devices; SKU: UCT-2112-198). Overall appearance was assessed using a clinical scoring matrix assigned after assessing the following parameters: significant (10-24%) body weight loss, neurological signs (hind limb paralysis, weak grip, ataxia), appearance (ruffled fur, hunched), responsiveness (low to moderate), and behavior (circling, head tilt). Mice that scored higher than three on two consecutive days were euthanized and documented as dead for the following day.

### Tissue collection

At two days postinfection, mice were anesthetized using 1–3% isoflurane, followed by euthanasia using an overdose of ketamine/xylazine. Whole blood (200 µl) was collected from the heart of mice and transferred into Eppendorf tubes. Serum was collected from blood by centrifugation at 3500 RPM and 4°C for 10 min and stored at -80°C for later analysis. 20-30 mg of liver, small intestine, brain, spleen, and kidney tissue were weighed and transferred into Eppendorf tubes holding 600 µL RNAlater (Ambion) and stored at 4°C for next day processing or at -80°C for future processing.

### RNA extraction from serum and tissues

The isolated tissues (20 to 30 mg) were transferred into 2 mL tubes containing buffer RLT–1 % β-mercaptoethanol (Qiagen) and a 5 mm stainless steel bead. Tissues were lysed using a TissueLyser (Qiagen) at 30 cycles/s for 2 min, 1 min wait, 30 cycles/s for 2 min, and centrifuged at high speed (13,000 rpm) for 10 min. Total RNA was then extracted from the resulting supernatant or serum using a RNeasy minikit (Qiagen) following the manufacturer’s instructions.

### Sera titrations

Viral particle quantification in sera was performed by plaque assay on HuH7.5 cells. 4.5×10^4^ cells were plated in 48-well plates 16-24 hours prior to experiments. For titration, sera were diluted 1:10 in OptiMEM + GlutaMax (Gibco) then serial diluted (10^-1^ – 10^-8^). 100 μL of each dilution was plated onto HuH7.5 cells, incubated for 2 hours at 37°C and 5% CO_2_, then the inoculum was removed prior to overlay of 800 μμL of 1% Methylcellulose media in DMEM 10% FBS. Plates were incubated for 5 days at 37°C and 5% CO_2_ before removal of overlay and fixation with 10% neutral-buffered formalin. After 2 hours of fixation, formalin was removed, and the fixed cells were stained with 0.1% crystal violet in 10% ethanol for 30 minutes. Cells were then washed and plaques were counted.

### Data representation and statistical analyses

Data are presented and analyzed using GraphPad Prism software v10.0.2. Significance was calculated as indicated in the corresponding figure legend.

## Supplementary Materials

Fig.S1. IFI6 does not interact with individual TBEV proteins.

Table S1. siRNA sequences used for knockdown experiments.

Table S2. RT-qPCR primer sequences.

Table S3. sgRNA sequences used for lentiviral vector-mediated CRISPR/Cas9 KO.

## Acknowledgments

We thank Sergei Kotenk at Rutgers University (NJ, USA), N. Escriou (Institut Pasteur, Paris, France), Nick Johnson (Animal and Plant Health Agency, Addlestone, Surrey, UK), Nolwenn Dheilly (Institut Pasteur, Paris, France), Gert Zimmer (Institute of Virology and Immunology (IVI), Bern, Switzerland), Olivier Schwartz (Institut Pasteur, Paris, France), and Nathalie Morel for providing reagents. FS was supported by a Boehringer Ingelheim Fonds PhD fellowship to realize this work.

---

**Fig. S1.**
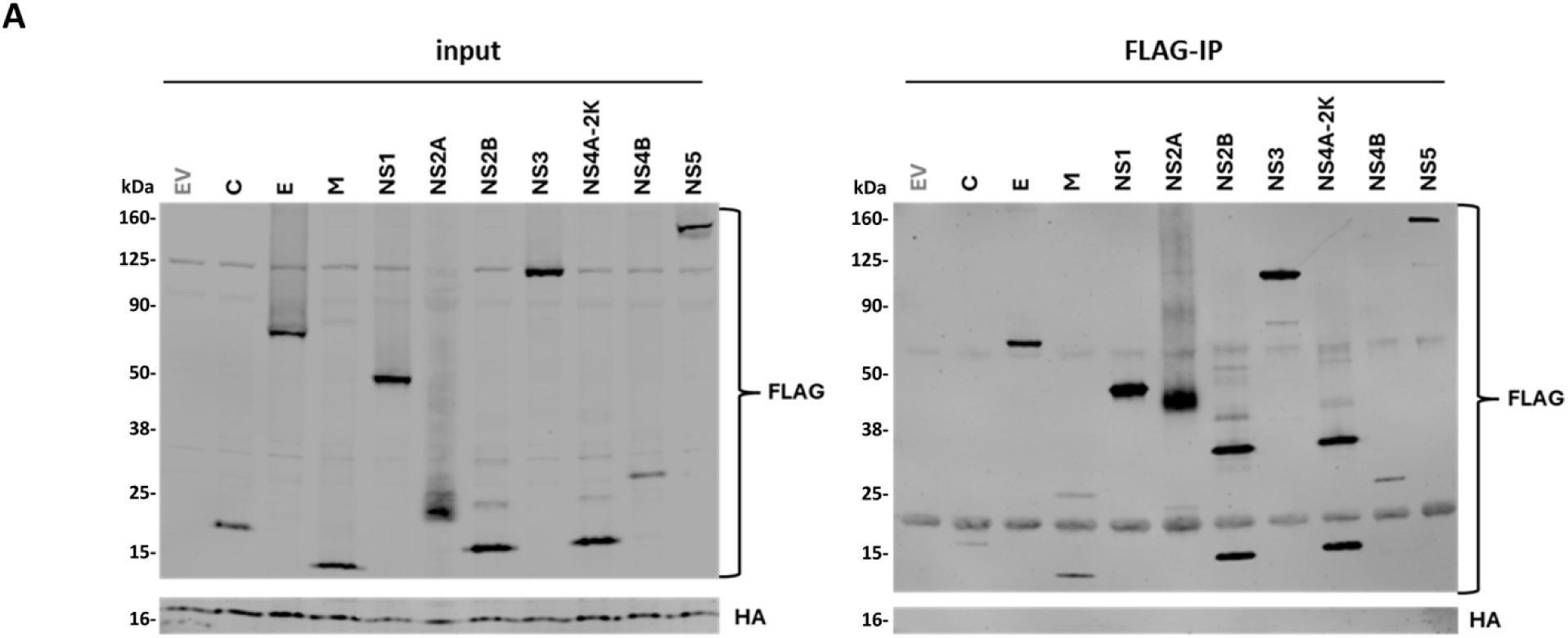
IFI6 does not interact with individual TBEV proteins. 293T cells were transfected with plasmids expressing IFI6-HA and empty vector (EV) plasmids or plasmids expressing FLAG tagged versions of proteins encoded by TBEV. Cells were lysed 24 h post transfection, and whole cell lysates were immunoblotted with the indicated antibodies (input). The same samples were issued to an immunoprecipitation assay using anti FLAG magnetic beads and subsequent staining with the indicated antibodies (FLAG-IP) (n=2).

**Table S1.**
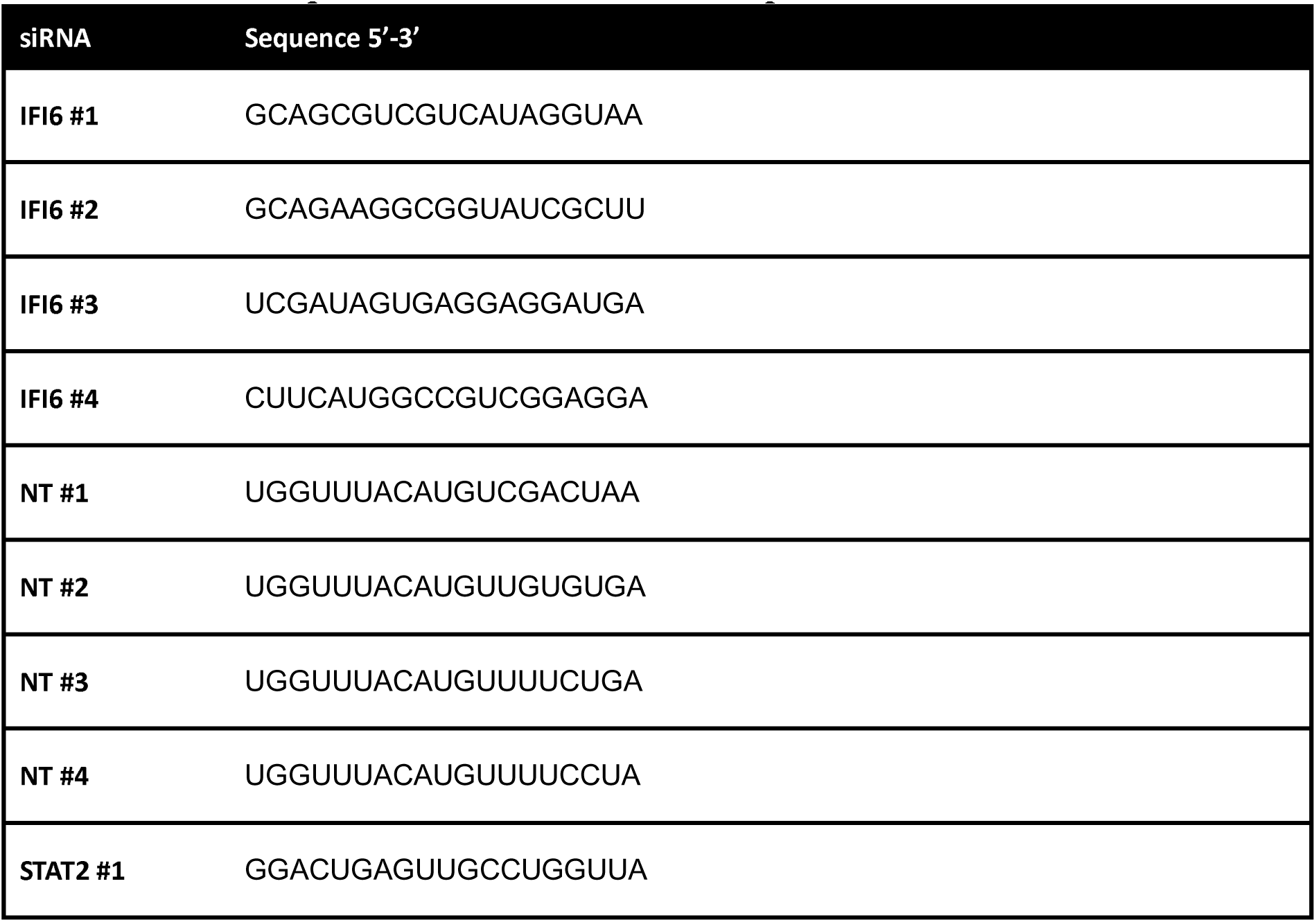
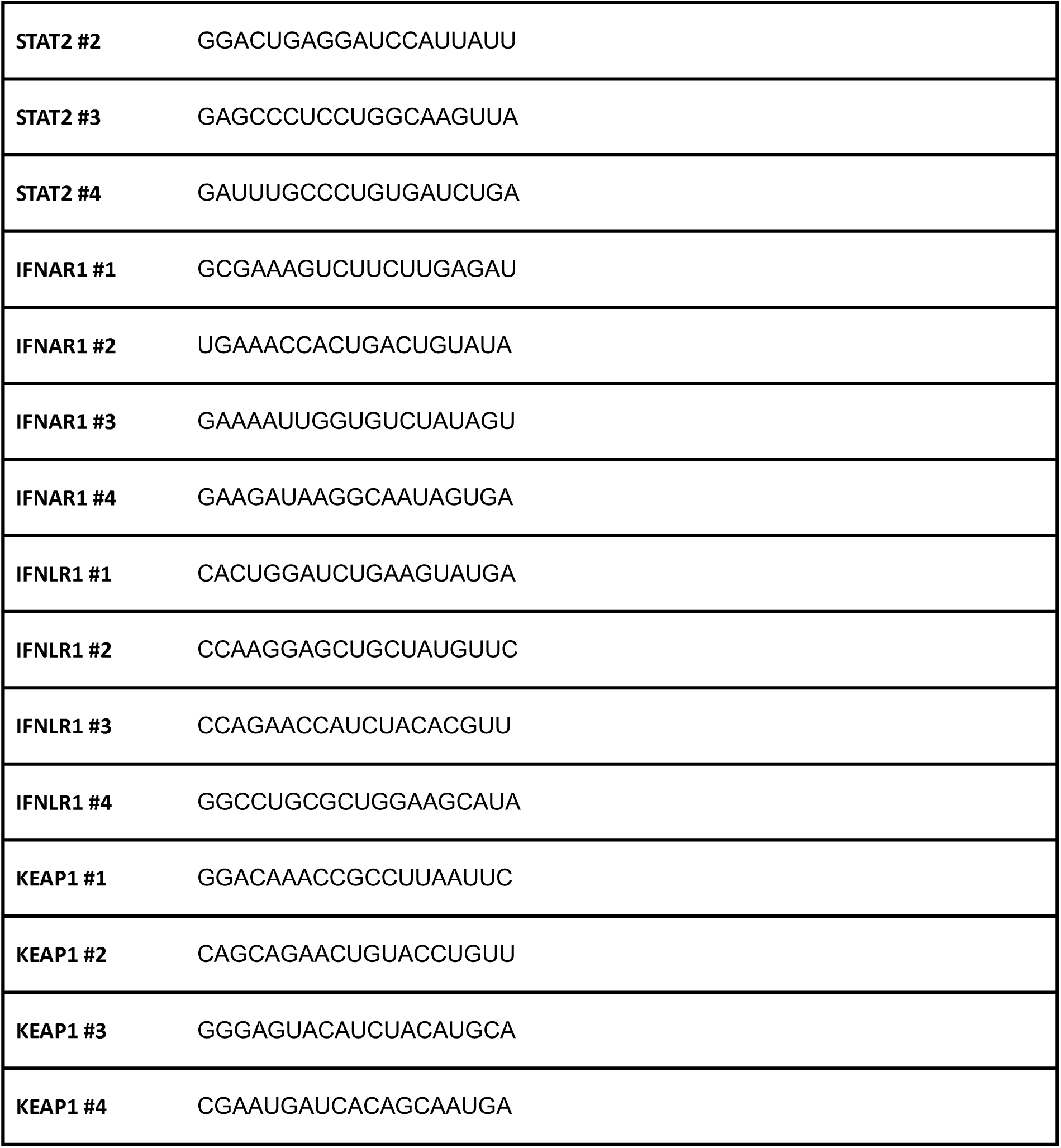
siRNA sequences used for knockdown experiments.

**Table S2.**
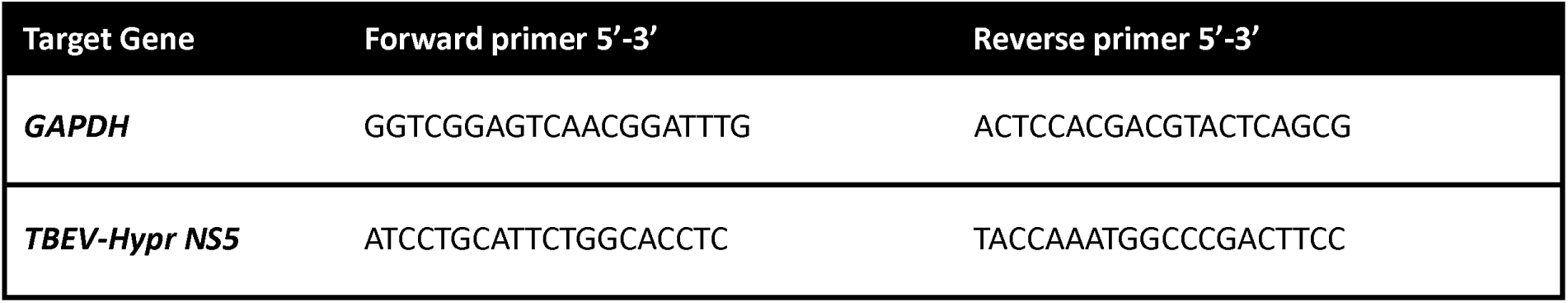
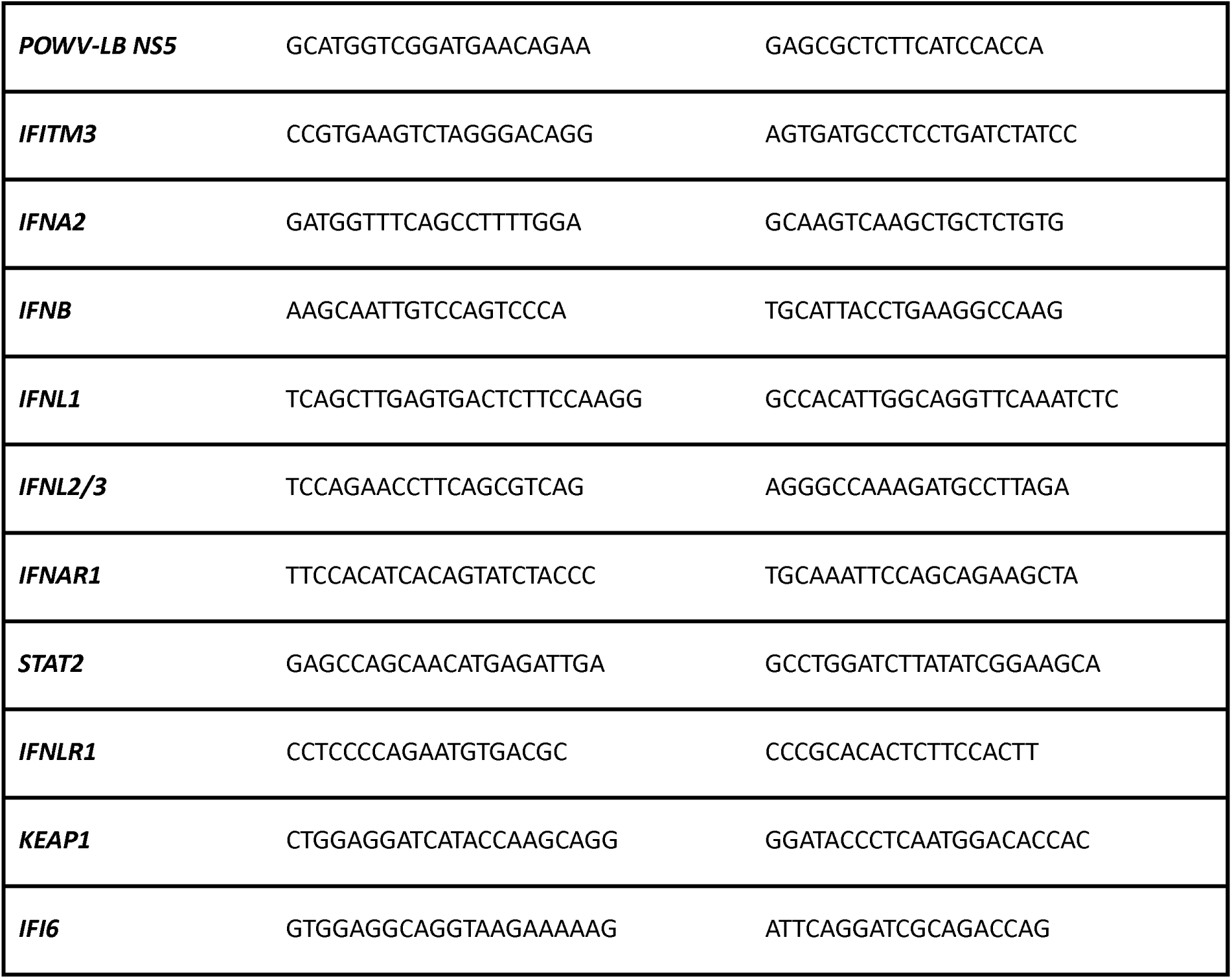
RT-qPCR primer sequences.

**Table S3.**
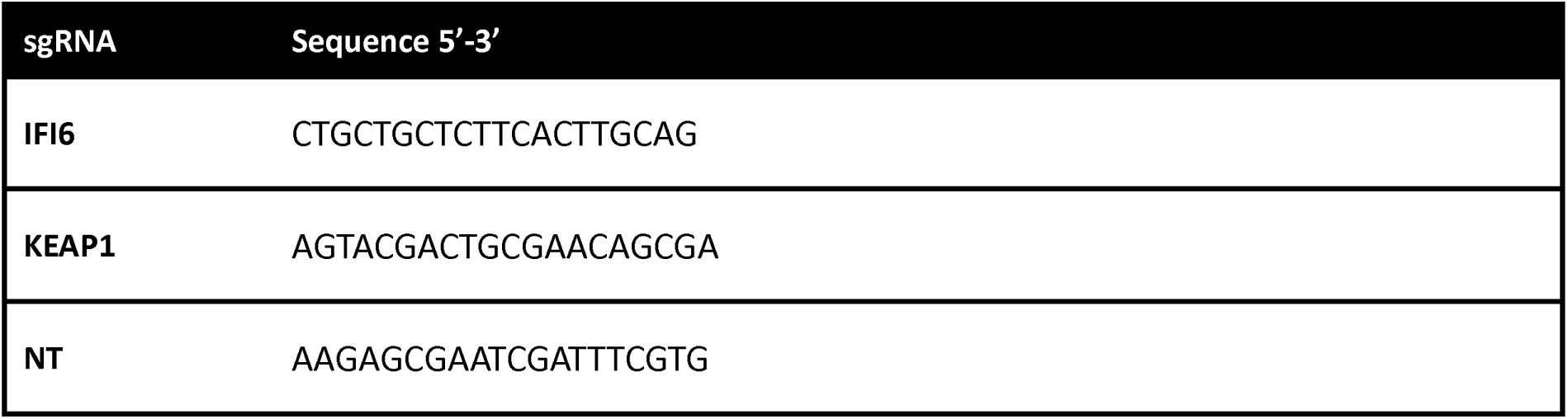
sgRNA sequences used for lentiviral vector-mediated CRISPR/Cas9 KO.

